# Epigenome-wide profiling in the dorsal raphe nucleus highlights cell-type-specific changes in *TNXB* in Alzheimer’s disease

**DOI:** 10.1101/2023.08.28.555168

**Authors:** RJM Riemens, E Pishva, A Iatrou, J Roubroeks, J Nolz, R Lardenoije, M Ali, A Del Sol, R Delgado-Morales, M Esteller, G Kenis, BPF Rutten, KP Lesch, SD Ginsberg, P Coleman, J Mill, D Mastroeni, A Ramirez, T Haaf, K Lunnon, DLA van den Hove

## Abstract

Recent studies have demonstrated that the dorsal raphe nucleus (DRN) is among the first brain regions affected in Alzheimer’s disease. Hence, in this study we conducted the first comprehensive epigenetic analysis of the DRN in AD, targeting both bulk tissue and single isolated cells. The Illumina Infinium MethylationEPIC BeadChip array was used to analyze the bulk tissue, assessing differentially modified positions (DMoPs) and regions (DMoRs) associated with Braak stage. The strongest Braak stage-associated DMoR in *TNXB* was targeted in a second patient cohort utilizing single laser-capture microdissected serotonin-positive (5-HT+) and -negative (5-HT-) cells isolated from the DRN. Our study revealed previously identified epigenetic loci, including *TNXB* and *PGLYRP1*, and novel loci, including *RBMXL2*, *CAST*, *GNAT1*, *MALAT1*, and *DNAJB13*. Strikingly, we found that the methylation profile of *TNXB* depends both on disease phenotype and cell type analyzed, emphasizing the significance of single cell(-type) neuroepigenetic studies in AD.

## Introduction

Over the last years, prime sites of early interrogation of Alzheimer’s disease (AD) have been extended beyond brain regions such as the hippocampus and the entorhinal cortex. Recent studies have shown that the brainstem is among the first regions affected and that neurofibrillary tangles can already be observed in brainstem nuclei such as the dorsal raphe nuclei (DRN), even before the manifestation of clinical symptoms [1–4]. Moreover, it has been suggested that pathology spreads from the brainstem to subcortical areas, and, subsequently, to areas of the neocortex, marking the clinical stages of AD. The essential role of the brainstem in AD has further been supported by findings of magnetic resonance imaging, demonstrating volume reductions and structural deformations in this brain region [5, 6]. Further evidence supporting the involvement of the DRN in AD concerns the occurrence of non-cognitive behavioral and neuropsychological symptoms in AD, particularly in prodromal stages, including depression, and related disturbances in mood and emotions, as well as changes in appetite, respiration and circadian rhythm [2].

Research aimed at exploring the etiopathophysiology of AD has furthermore implicated a crucial role for epigenetic mechanisms, next to both independent and interdependent genetic, environmental, and lifestyle factors in the development and course of this disease [7, 8]. Accordingly, several recent epigenome-wide association studies (EWAS) in brain tissue derived from AD patients have identified differences in DNA methylation and hydroxymethylation profiles in the AD cortex across independent patient cohorts [9–21].

Despite the increasing number of EWAS in AD, examining the DRN for potential disease-specific deviant epigenetic signatures, indicative of the more incipient stages of the disease, has not been performed up to now. Additionally, there is a growing need for single-cell approaches for studying epigenetic dysregulation in AD [22], yet no studies applying such techniques for this disorder have been performed to date. Therefore, in this study, we performed the first EWAS of the DRN in AD. First, using DRN bulk tissue, we examined the association between Braak stage and alterations in three states of cytosine-phosphate-guanine (CpG) sites, including 5-methylcytosine (5-mC; reflecting DNA methylation), 5-hydroxymethylcytosine (5-hmC; reflecting DNA hydroxymethylation), and unmodified cytosine (5-uC; reflecting positions without any type of modification). Following validation of the EWAS discovery findings using bisulfite pyrosequencing in the same cohort, we utilized single-cell, limiting dilution bisulfite pyrosequencing (LDBSP) to explore the cell-specific nature of the DRN’s epigenetic signature. This investigation was conducted in an independent patient cohort, targeting bisulfite methylation patterns in both serotonin (5-hydroxytryptamine [5-HT])-positive (5-HT+) and non-serotonergic (5-HT-) cells, isolated from the DRN through laser capture microdissection (LCM).

## Results

### DMoPs associated with Braak stage in the DRN bulk tissue

Epigenetic profiles at the level of 5-uC, 5-mC and 5-hmC were quantified over 850.000 single CpG sites using Illumina Infinium MethylationEPIC BeadChip arrays and DNA isolated from DRN bulk tissue. Samples of 93 donors were collected from the MRC London Brain bank for Neurodegenerative Disease (see Table 1 for the cohorts’ demographics). Our discovery cohort represented the entire spectrum of AD pathology defined by Braak stage, hence we focused on identifying Braak stage-associated differentially modified positions (DMoPs) at the level of all three CpG modifications, *i.e.* differentially unmodified positions (DUPs), differentially methylated positions (DMPs), and differentially hydroxymethylated positions (DHPs), for 5-uC, 5-mC, and 5-hmC, respectively. None of the DMoPs identified reached the false discovery rate (FDR) threshold for genome-wide significance. Probes with nominal *p* values <1.0E-3 were ordered based on a combined *p* value and regression estimate ranking. The summary of statistics related to the top-ranked nominally significant DUPs (1029 probes), DMPs (647 probes) and DHPs (831 probes) are shown in Supplementary Tables 1-3, respectively.

**Table 1.** Demographics of studied cohorts.

|  | **EWAS**<br>**(Discovery)** | **Pyrosequencing**<br>**(Technical validation)** | **LDBSP and pyrosequencing**<br>**(Cell-specific validation)** |  |
| --- | --- | --- | --- | --- |
| **Tissue or cell-type** | DRN bulk tissue | DRN bulk tissue | 5-HT+ cells | 5-HT- cells |
| **N** | 93 | 62 | 24 (12/12; AD/C) | 20 (10/10; AD/C) |
| **Sex (F/M)** | 53/40 | 33/29 | 24/0 | 20/0 |
| **Age of death; Mean (± SD)** | 83.85 (9.04) | 84.78 (9.14) | 81.21 (10.21) | 81.35 (11.31) |
| **PMI; Mean (± SD)** | 32.66 (20.04) | 43.35 (89.05) | 2.69 (0.82) | 2.77 (0.87) |
| **Braak Stage; Range (Median)** | 0-VI (V) | 0-VI (V) | I-VI (IV) | I-VI (IV) |

Displayed are the demographics of the patient cohorts that were used in the present study. Dorsal raphe nuclei (DRN) tissues used in the discovery epigenome-wide association study (EWAS) and pyrosequencing (technical) validation study were obtained from the Medical Research Council (MRC) London Brainbank for Neurodegenerative Disease (London, UK). Serotonergic and non-serotonergic cells, i.e. serotonin-positive (5-HT+) and 5-HT-negative (5-HT-) cells, isolated from the DRN by means of laser capture microdissection (LCM) used in the cell-specific validation study were obtained from the Banner Sun Health Research Institute (Sun City, Arizona, USA). For all cohorts, the type of tissue or cells, the number of samples in each group, the distributions of sex (F, female; M, male), the age of death (Mean ± standard deviation (SD)), the postmortem interval (PMI; in hours) and the Braak stage are shown.

### DMoRs associated with Braak stage in the DRN bulk tissue

Next, we investigated the spatial correlation of adjacent modified CpG sites to identify differentially modified regions (DMoRs) in the DRN bulk tissue, which were defined as regions containing three or more correlated DMoPs that displayed a *p*_Šidák_ value <0.05 within a 500-bp sliding window. Hence, this analysis aimed at identifying genomic regions where the degree of Braak stage-associated change was significantly correlated between neighboring CpG sites at the level of all three modifications, independent of the directionality of change for these modifications, observed at the respective positions. In total, eight significant DMoRs were identified that survived multiple testing correction, which are listed in Table 2 and ranked based on their *p*_Šidák_ value. An overview of the DMoPs belonging to each of the DMoRs and their associated changes in this regional analysis, as well as the regression estimates (REs), the *p* values for each probe for the 5-uC, 5-mC or 5-hmC levels, and the bisulfite signal, *i.e.* 5-mC + 5-hmC, can be found in Supplementary Table 4.

**Table 2.** Differentially modified regions (DMoRs) in the dorsal raphe nuclei (DRN) bulk tissue.

| Rank | USCS annotation | Chromosome | Start | End | Num. of probes (modification) | *p* value | *p* Šidák value | Gene feature | CpG island feature |
| --- | --- | --- | --- | --- | --- | --- | --- | --- | --- |
| 1 | TNXB | 6 | 32063459 | 32064033 | 15 (14, 5-uC; 1, 5-mC) | 7,25E-15 | 9,45E-12 | CDS | Island |
| 2 | PGLYRP1 | 19 | 46526321 | 46526652 | 9 (9, 5-uC) | 7,91E-13 | 1,80E-09 | Intergenic | Island |
| 3 | RBMXL2 | 11 | 7110074 | 7110149 | 5 (4, 5-uC; 1, 5-mC) | 3,18E-09 | 3,18E-05 | Intergenic | Island |
| 4 | CAST | 5 | 95997186 | 95997356 | 4 (4, 5-uC) | 4,50E-09 | 1,99E-05 | Intergenic | Shore |
| 5 | GNAT1 | 3 | 50230792 | 50230886 | 3 (3, 5-uC) | 1,38E-08 | 1,10E-04 | Intron + CDS | Island |
| 6 | MALAT1 | 11 | 65266482 | 65266562 | 4 (4, 5-mC) | 1,619E-08 | 1,52E-04 | NC exon | Shore |
| 7 | DNAJB13 | 11 | 73668626 | 73668762 | 3 (1, 5-uC; 2, 5-mC) | 2,966E-08 | 1,64E-04 | Intron; Intron + 5'UTR | NA |
| 8 | TNXB | 6 | 32064785 | 32064957 | 6 (6, 5-uC) | 2,094E-07 | 9,14E-04 | CDS | Island |

For each DMoR in the DRN bulk tissue, the Illumina gene annotation (UCSC Annotation), the chromosomal position and coordinates (Ensembl GRCh37 assembly), the number of probes, the *p* value, the multiple testing corrected *p* Šidák value, the gene feature (CDS, coding sequence; NC, non-coding; 5'UTR, 5'untranslated region) and the cytosine-phosphate-guanine (CpG) island feature are shown. All regions are ranked based on their *p* Šidák value with *p* Šidák < 0.05 as a cut-off.

The most significant DMoR (*p*_Šidák_ = 9.45E-12) was identified in the coding DNA sequence of *TNXB,* demonstrating DNA demethylation in this locus with advancing AD neuropathology. The annotated region spanned 575 bp (6:32063459-32064033) and contained fifteen probes, of which fourteen were related to an increase in 5-uC, including cg14196170 (chr6:32063595, RE = 14.72, *p* = 2.93E-05) and cg27387193 (chr6:32064032, RE = 11.28, *p* = 2.76E-04) that were also listed among the top-ranked DUPs, whereas one probe, cg18460422, demonstrated a decrease in 5-mC.

The next significant hit (*p*_Šidák_ = 1.80E-09) was found in an intergenic region of *PGLYRP1*, displaying a positive relationship between 5-uC and disease progression for each regionally associated probe, hence demonstrating that an overall loss of DNA modifications is typically observed in AD for this locus. The DModR in this gene spanned 332 bp and covered nine probes (chr19:46526321-46526652), of which cg08682981 (chr19:46526489, RE = 33.43, *p* = 1.7E-06), cg14131693 (chr19:46526651, RE = 28.46, *p* = 2.56E-05), and cg25754173 (chr19:46526453, RE = 26.19, *p* = 5.49E-04), were also listed among the top-ranked DUPs.

A similar Braak stage-associated loss of DNA modifications was observed for the subsequent three significant hits. For the DMoR in the intergenic region of *RBMXL2* (*p*_Šidák_ = 3.18E-05), consisting of 5 probes and spanning 76 bp (chr11:7110074-7110149), an increase for 5-uC in four out of the five sites and a decrease of 5-mC for the other probe, was identified. Two out of these four 5-uC probes, *i.e.* cg23916104 (chr11:7110083, RE = 5.20, *p* = 4.83E-04) and cg21949858 (chr11:7110145, RE = 3.43, *p* = 8.20E-04), were also listed among the top-ranked DUPs. For the four-probe long DMoR in *CAST*, which spanned 171 bp (chr5:95997186-95997356; *p*_Šidák_ = 1.99E-05), and included cg16381797 (chr5:95997312, RE = 55.52, *p* = 6.30E-05) and cg14097306 (chr5:95997355, RE = 13.34, *p* = 2.95E-04) from the top-ranked DUPs, a positive association with Braak stage at the level of 5-uC was identified.

In addition, the 95 bp region in *GNAT1* (chr3:50230792-50230886; *p*_Šidák_ = 1.10E-04), included three probes, all showing a Braak stage-associated increase for 5-uC, and of which cg23359897 was listed both among the top-ranked DUPs (chr3:50230792, RE = 30.51, *p* = 1.49E-06) and DMPs (chr3:50230792, RE = −10.55, *p* = 2.08E-04).

Another region annotated to *MALAT1* containing 4 probes and spanning 81 bp (chr11:65266482-65266562) also showed up as significant DMoR (*p*_Šidák_ = 1.52E-04). A positive association was observed for 5-mC for each of these latter CpG sites, demonstrating that an increase in DNA methylation in this locus is associated with increasing neuropathology. From these probes, three were listed among the top-ranked DMPs, that is cg04378603 (chr11:65266494, RE = 47.83, *p* = 1.65E-04), cg05491695 (chr11:65266512, RE = 36.91, *p* = 2.93E-04), and cg04977124 (chr11:65266482, RE = 21.55, *p* = 2.85E-04). This was the only gene showing a Braak stage-associated increase in DNA methylation, since for the remaining two DMoRs also an overall loss of CpG modifications was observed.

The DMoR in *DNAJB13,* consisted of 80 bp (chr11: 73668626-73668762; *p*_Šidák_ = 1.64E-04) and contained 3 probes, of which cg20737388 and cg04044120 demonstrated changes in 5-mC and cg04950148 in 5-uC. While a negative relationship was apparent for the 5-mC probes, a positive association with disease progression was identified for the 5-uC probe. Furthermore, all three CpG sites were listed among the top-ranked hits, of which cg04044120 (chr11:73668757, DUP, RE = 10.92, *p* = 7.07E-04; DMP, RE = −7.052, *p* = 1.64E-04) was part of both the DUPs and DMPs, and cg04950148 (chr11:73668761, DUP, RE = 9.78, *p* = 8.63E-04) and cg20737388 (chr11: 73668626, DMP, RE = - 4.450226021, *p* = 5.40E-04), of the DUPs and DMPs, respectively. Finally, our last FDR-corrected significant hit represented a second DMoR in *TNXB* more downstream of the first most significant Braak stage-associated region, covering 172 bp (6:32064785-32064957; *p*_Šidák_ = 9.14E-04), for which an increase in 5-uC was observed at all six CpG sites with advancing neuropathology.

In parallel to the novel DMoR analysis described above, we also conducted the classical regional analyses previously performed in AD EWAS, which is aimed at identifying regions containing either three or more correlated DUPs, DMPs, or DHPs, that displayed a *p*_Šidák_ value <0.05 within a 500-bp sliding window, *i.e.* differentially unmodified regions (DURs), differentially methylated regions (DMRs), and differentially hydroxymethylated regions (DHRs), respectively. Hence, genomic regions were identified where the degree of Braak stage-associated change was significantly correlated between neighboring CpG sites at the level of only one type of cytosine modification. The results of these analyses are listed in Supplementary Table 5 and ranked based on their *p*_Šidák_ value. Five significant DURs in *PGLYRP1*, *TNXB*, *CAST*, *GNAT1*, and *RBMXL2*, and one DMR in *MALAT1*, that survived FDR correction, were identified. As expected, the identified regions in the classical regional analyses largely overlapped with the identified DMoRs. Remarkably, our DMoR analysis was able to identify two additional regions in *TNXB* and *DNAJB13*, which could not be obtained with the classical approach for identifying Braak stage-associated regions. Thus, these latter findings highlight the informative nature of the DMoR approach proposed in this study, demonstrating superior capability in detecting dysregulated epigenetic loci across all three cytosine states (5-uC, 5-mC, and 5-hmC).

### Structural and functional genomic annotation enrichment analysis based on DMoPs

We then assessed biological, cellular and molecular pathways in the DRN bulk tissue that were enriched among each of the Braak stage-associated DMoPs. For this purpose, a Gene Ontology (GO) enrichment analysis using the top-ranked DUPs, DMPs and DHPs was performed (Supplementary Tables 6-8). We identified two pathways related to the DHPs that surpassed the FDR threshold for multiple testing correction (FDR-adjusted *p* <0.05). The most significant GO term in our 5-hmC enrichment analyses was found in ‘homophilic cell adhesion via plasma membrane adhesion molecules’ (FDR adjusted *p* = 1.16E-03), demonstrating that from the 167 annotated genes, 27 were differentially modified. The second significantly enriched GO term that was associated with our 5-hmC data was found for ‘calcium ion binding’ (FDR-adjusted *p* = 1.09E-02), revealing that from the 690 genes, 56 were altered.

### Technical validation of changes in *TNXB* DNA methylation in the DRN bulk tissue using bisulfite pyrosequencing

To validate our discovery findings, we then compared our array-based data to bisulfite pyrosequencing data generated from a subset of identical DNA samples (n = 62) obtained from the DRN bulk tissue samples (Table 1). Importantly, bisulfite pyrosequencing does not allow for the segregation of 5-mC and 5-hmC [22]. Therefore, we aimed at validating the most significant Braak stage-associated DMoR in *TNXB* by assessing the level of bisulfite methylation, *i.e.* the cumulative measure of 5-mC and 5-hmC, which is inversely proportional to the measurement of 5-uC [23]. Of note, an overlapping region in *TNXB* when compared to the DMoR and DUR analyses was also significantly associated with Braak stage when performing the regional analyses for the bisulfite dataset only (See Supplementary Table 9; *TNXB*, chr6:32063595-32064033, *p*_Šidák_ = 5,71E-09). The probes annotated to this differentially bisulfite methylated region (DbsMR), all demonstrated a negative association with Braak stage, i.e. opposite to the association at the level of 5-uC for the annotated probes in the DMoR and DUR of this locus. These various regional analyses collectively indicated that an overall increase in 5-uC and a decrease in 5-mC plus 5-hmC in *TNXB* is associated with progressing AD neuropathology.

The DNA bisulfite methylation status of *TNXB* in the samples belonging to our technical validation cohort, was quantified across a region of 72 bp (chr6:32063869-32063940), spanning eight CpG sites. Hence, the targeted region overlapped with the most significant DMoR (Figure 1), the DUR, as well as the DbsMR, including three Illumina probes, *i.e.* cg10365886, cg14188106 and cg07524919. The results from the linear regression analysis on the *TNXB* bisulfite pyrosequencing data are provided in Table 3, whereas the results from the Pearson correlation analyses between the EWAS-derived bisulfite data set and the bisulfite pyrosequencing data are shown in Figure 2. The regression analysis revealed a strong significant association with Braak stage for all eight targeted CpG sites, that is all *p* values were <1.73E-03, with the smallest *p =* 1.63E-04 for CpG site four (cg07524919) in the sequence. In addition, the correlation analysis revealed that the pattern of DNA bisulfite methylation in *TNXB* quantified using the BeadChip arrays was identical to the bisulfite pyrosequencing data, with a highly significant and strong correlation between the values estimated by the two independent technologies for all three analyzed Illumina probes. The first CpG site in the sequenced region (cg10365886) displayed the smallest *p*-value of 2.2E-16 and a correlation coefficient of 0.84.

**Figure 1.**
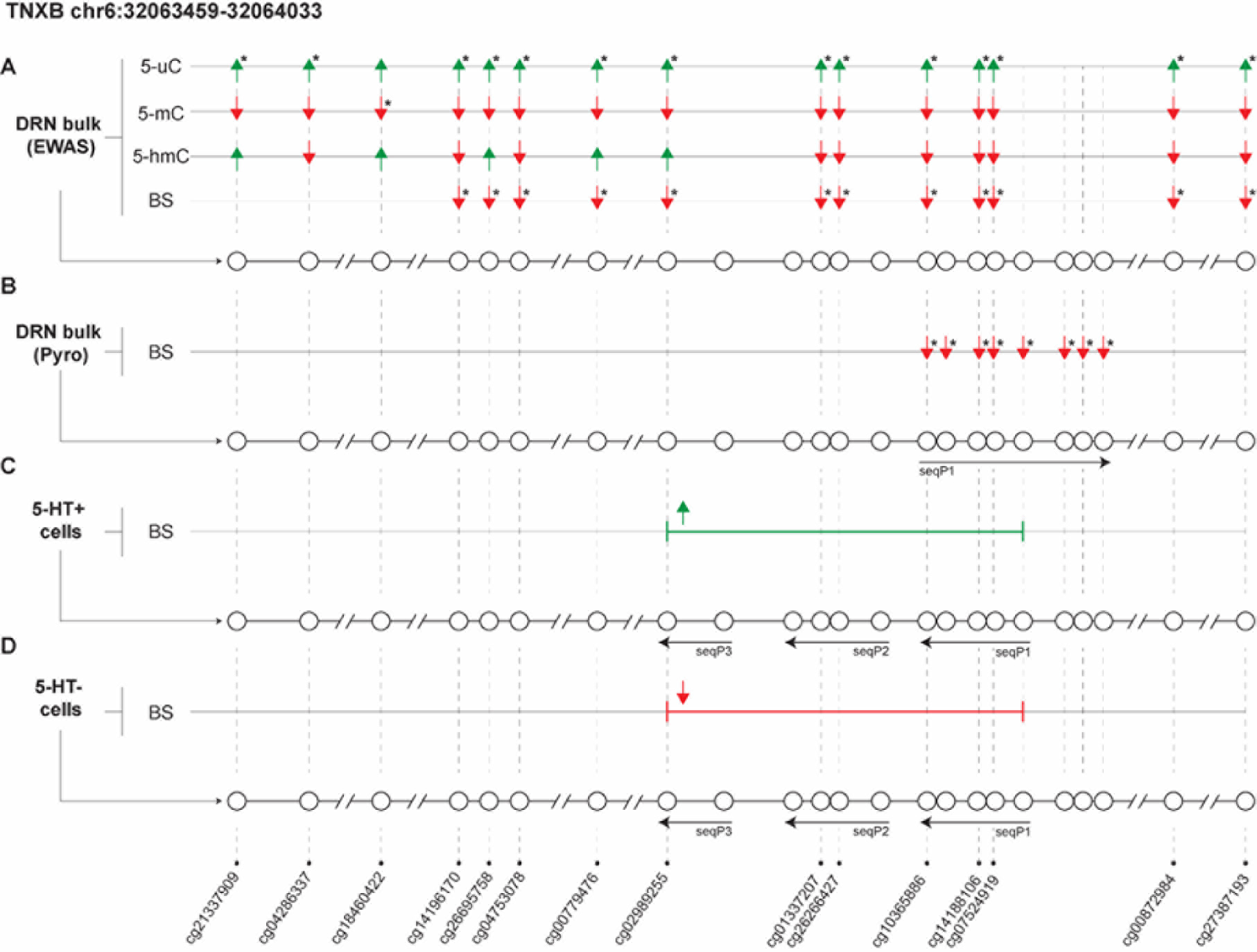
Overview of the Alzheimer’s disease (AD)-associated region in Tenascin XB (*TNXB*) in the dorsal raphe nucleus (DRN). **A**. Shown for the DRN epigenome wide association study (EWAS) is the differentially modified region (DMoR; chromosome(chr)6:32063459-32064033) based on three cytosine states, *i.e.* unmodified cytosine (5-uC), methylated cytosine (5-mC), and hydroxymethylated cytosine (5-hmC). For each cytosine-phosphate-guanine (CpG) site, the direction of change for all of the aforementioned modifications, based on the individual CpG regression estimates (REs), are depicted by and arrow, where ↑ means an increase and ↓ a decrease associated with Braak stage. CpG sites and their respective modification belonging to the significant DMoR are indicated with an asterisk. In addition, the differentially bisulfite methylated region (DbsMR) in *TNXB* that is based on the bisulfite (BS) signal only, representing the cumulative measure of 5-mC and 5-hmC, is depicted. Similarly, the direction of change for each DbsMR-associated CpG site is indicated with an arrow and CpG sites belonging to the significant DbsMR are indicated with an asterisk. **B.** Shown for the technical validation analysis in the DRN bulk tissue using BS pyrosequencing (Pyro) are the targeted CpG sites in the DMoR of *TNXB*. The direction of change associated with Braak-stage based on the individual REs are indicated with an arrow and the asterisks indicate all CpG sites that were significant during the regression analysis. The horizontal arrow indicates the orientation of the pyrosequencing assay that was based on one sequencing primer (seqP). **C.** For the cell type-specific validation analysis in the serotonergic, *i.e.* serotonin-positive (5-HT+) cells, the region that was targeted in the DMoR of *TNXB* is shown. The average change in CpG BS methylation levels associated with AD is depicted with an arrow. The horizontal arrows indicate the orientation of the pyrosequencing assay that was based on three seqPs. **D.** Similarly, for the non-serotonergic, *i.e.* 5-HT-negative (5HT-) cells, the targeted region and the average disease-associated change in CpG BS methylation levels is shown.

**Figure 2.**
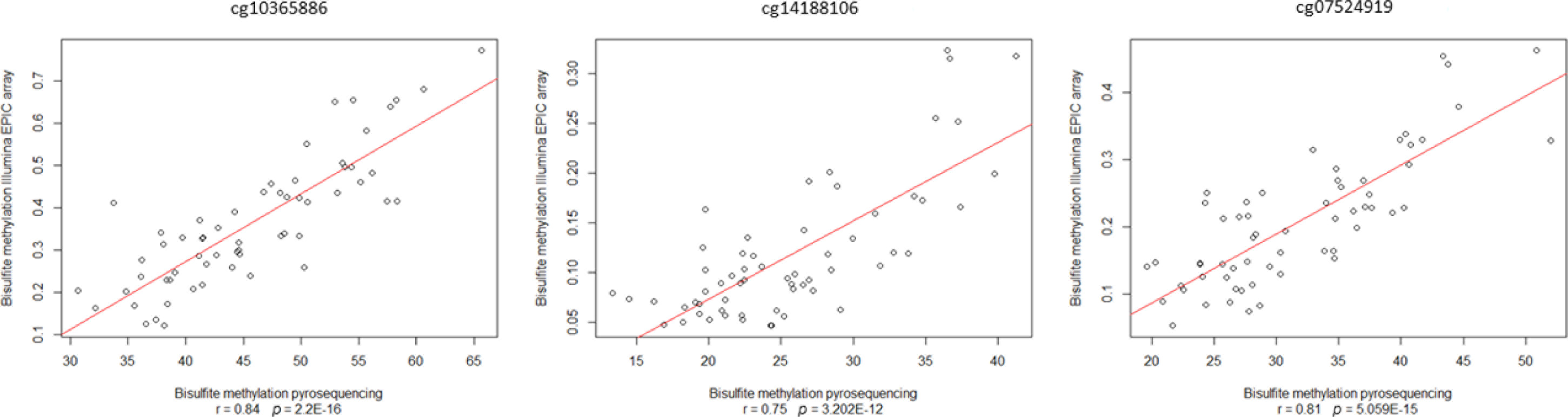
Correlation analysis during the validation study of the Tenascin XB (*TNXB)* Braak-associated region in the dorsal raphe nucleus (DRN) bulk tissue. DNA bisulfite methylation patterns quantified by the Illumina Infinium MethylationEPIC BeadChip array and bisulfite pyrosequencing in *TNXB* were highly significantly and strongly correlated for cg10365886, cg14188106 and cg07524919.

**Table 3.** Regression analysis during the validation of the *TNXB* Braak-associated region in the dorsal raphe nuclei (DRN) bulk tissue.

| **CpG** | **Probe** | **RE** | **SE** | **t value** | **p-value** |
| --- | --- | --- | --- | --- | --- |
| 1 | cg10365886 | -0,10599 | 0,02909 | -3.644 | 5,75E-04 *** |
| 2 | - | -0,10048 | 0,02656 | -3.783 | 3,69E-04 *** |
| 3 | cg14188106 | -0,12461 | 0,03575 | -3.486 | 9,43E-04 *** |
| 4 | cg07524919 | -0,12121 | 0,03006 | -4.032 | 1,63E-04 *** |
| 5 | - | -0,12586 | 0,03256 | -3.866 | 2,82E-04 *** |
| 6 | - | -0,09968 | 0,03033 | -3.286 | 1,73E-03 ** |
| 7 | - | -0,12871 | 0,03687 | -3.491 | 9,36E-04 *** |
| 8 | - | -0,12202 | 0,03827 | -3.188 | 2,34E-03 ** |

The DNA bisulfite methylation patterns quantified by pyrosequencing over eight cytosine-phosphate-guanine (CpG) sites in *TNXB*, including cg10365886, cg14188106 and cg07524919, were all significantly associated with Braak stage in the DRN. Displayed for each CpG site is their respective number in the sequence, the Illumina probe annotation (if applicable), the regression estimate (RE) for the Braak stage-associated analysis, the standard error (SE), the *t*-statistics and accompanying p values. \*\* = $p < 0.01$ ; \*\*\* = $p < 0.001$

Thus, based on these data, we were able to robustly validate our EWAS discovery findings in the Braak stage-associated region of *TNXB*. Moreover, in addition to the analyzed Illumina probes, we also demonstrated that adjacent CpG sites in the region of this gene that were not covered on the Illumina array, support the notion of an overall decrease in 5-mC and 5-hmC with increasing Braak stage. Altogether, these data provide compelling evidence for an association between active DNA demethylation in this *TNXB* locus and an increase of AD-related neuropathological features.

### Cell type-specific validation analysis in serotonergic and non-serotonergic cells in the DRN

To validate our previous findings in an independent patient cohort obtained from the Banner Sun Health Research Institute (BSHRI) Brain and Body Donation Program (BBDP) [24], we first employed laser capture microdissection (LCM) to isolate individual DRN cells, followed by a novel cell type-specific and targeted epigenetic analysis called Limiting Dilution Bisulfite Pyrosequencing (LDBSP). This targeted approach was performed with the additional aim to define the cellular origin of the identified epigenetic signature in *TNXB*. For this purpose, 150 5-HT+ cells were isolated per individual from post-mortem DRN tissue derived from 12 cases without (Control cases; Braak I-IV) and 12 cases with a clinical diagnosis of AD (AD cases; Braak V-VI) (Table 1). LDBSP was then applied on the 5-HT+ cells to analyze the bisulfite methylation status of the Braak stage-associated region in *TNXB* on single alleles [25].

The targeted region in this analysis consisted of 140 bp (chr6:32063774-32063913) and spanned eleven CpG sites (Figure 1), thereby overlapping with the most significant DMoR, the DUR, and the DbsMR, identified for *TNXB* in the DRN bulk tissue, as well as overlapping partially with the targeted region during our technical validation analysis. From the eleven CpG sites in this region, six represented Illumina probes that were also part of the Braak stage-associated regions, *i.e.* cg10365886, cg14188106, cg07524919, cg26266427, cg01337207 and cg02989255. From these, the first three probes were those targeted during the technical validation analysis.

The results of the LDBSP analysis in 5-HT+ cells using a case-control model based on clinical diagnosis of the disease can be found in Figure 3. The data that is shown represents the average bisulfite methylation levels measured over the analyzed CpG sites in these cell types. We observed an overall trend towards AD-associated bisulfite hypermethylation for the *TNXB* CpG sites in the 5-HT+ cells. Surprisingly, this pattern contradicted our previous findings from the conducted EWAS on DRN bulk tissue. Accordingly, we then hypothesized that the EWAS-associated *TNXB* hypomethylation should originate from other, non-serotonergic, cells in the DRN bulk tissue.

**Figure 3.**
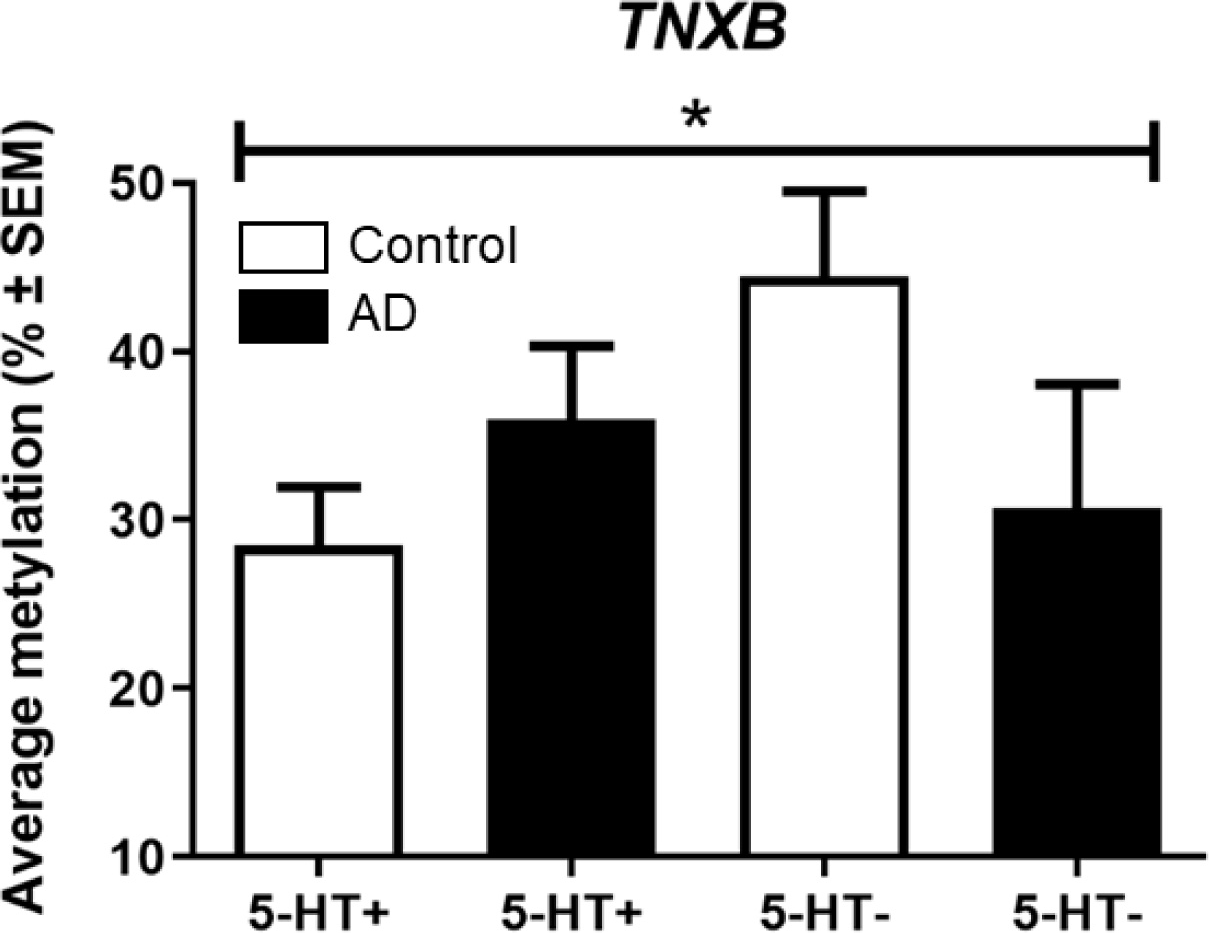
General linear model during the cell type-specific validation study in the dorsal raphe nuclei (DRN) of the Braak-associated region in Tenascin XB (*TNXB*). Average DNA bisulfite methylation levels quantified in both the serotonergic and non-serotonergic cells, *i.e.* serotonin-positive (5-HT+) and 5-HT-negative (5-HT-) cells, derived from the DRN of Alzheimer’s disease (AD) patients and healthy controls (Control) are shown. A general linear model with experimental condition as a between-subject factor and cell type as a within-subject factor revealed a significant interaction effect between cell type and experimental condition for *TNXB*. The AD-associated bisulfite methylation profiles were exactly opposite in the 5-HT+ cells when compared to the 5-HT-cells, the latter which resembled our EWAS data in the DRN. For the interaction effect: * = *p < 0.05*.

To confirm this hypothesis, we then isolated non-serotonergic, 5-HT-cells, by LCM from the exact same BBDP DRN tissue samples used in the LDBSP analysis. As such, regions that were free of 5-HT immunoreactivity, most likely consisting of non-serotonergic parenchymal cellular populations residing in the DRN, were isolated [26]. The average bisulfite methylation status in the exact same region of *TNXB* was then quantified in the 5-HT-cells, by using bisulfite pyrosequencing with an adapted pre-amplification protocol (see Materials and methods).

Subsequently, the average bisulfite methylation levels over all CpG sites in the targeted region of *TNXB* were then compared between both experimental groups and cell types using a general linear model with experimental condition (AD versus Control) as a between-subject factor and cell type (5-HT+ versus 5-HT-cells) as a within-subject factor. Accordingly, we found a significant interaction effect (*p* = 0.045) between experimental condition and cell type (Figure 3), with the average AD-associated bisulfite methylation profiles being exactly opposite in the 5-HT+ cells when compared to 5-HT-cells. More specifically, while the average CpG bisulfite methylation levels tended to be increased in 5-HT+ cells derived from AD patients, a decrease was observed in 5-HT-cells from the same patients when compared to healthy controls – the latter of which resembled the EWAS data in the DRN bulk tissue. Overall, these data obtained from our second independent patient cohort demonstrate, for the first time, that epigenetic dysregulation in *TNXB* in the DRN of AD patients is cell type-specific. Furthermore, these data corroborated our previous findings in the DRN discovery EWAS, indicating that AD-associated epigenetic dysregulation in *TNXB* likely reflects changes in non-serotonergic cells within this brainstem nucleus.

## Discussion

This study represents the first extensive epigenetic analysis performed in the DRN of AD to date. In our discovery cohort, an EWAS on bulk tissue derived from the DRN, was performed with the aim of discovering DMoPs associated with Braak stage at the level of three different CpG cytosine states, *i.e.* 5-uC, 5-mC and 5-hmC. In total, 1029, 647 and 831 nominally significant DUPs, DMPs DHPs, respectively, were identified for the aforementioned modifications, with a stringent *p* value cut-off of <1.0E-3. In a subsequent novel regional analysis looking at the spatial correlation of adjacent DMoPs, we then discovered eight significant FDR-adjusted DMoRs that were associated with Braak stage in the DRN bulk tissue. These included two regions in *TNXB*, and one in *PGLYRP1*, *RBMXL2*, *CAST*, *GNAT1*, *MALAT1*, and *DNAJB13*. We then successfully followed up the EWAS with a technical validation of our most significant DMoR in *TNXB* using bisulfite pyrosequencing in a subset of DNA samples derived from the exact same cohort. In addition, our DRN EWAS was complemented with a novel cell type-specific analysis, quantifying bisulfite methylation signatures in the previously identified region of *TNXB*, in individually isolated DRN 5-HT+ and 5-HT-cells derived from a second, independent patient cohort. As such, we aimed at validating our EWAS data and, for the first time to date, to simultaneously unravel potential cell type-specific contributions of the identified epigenetic signatures within the DRN.

Importantly, in this study, we performed a region-based analysis to identify the spatial correlation of neighboring CpG sites at the level of all cytosine states simultaneously and not for each of them (*i.e.*, 5-uC, 5-mC, and 5-hmC) individually, as performed in previously published EWAS in AD [14, 15, 21]. In fact, previous studies have been restricted to correlating an individual layer of CpG modifications or to correlating the CpG signatures derived from the bisulfite signal only, *i.e.* the combined signal of 5-mC and 5-hmC, with respect to a single measure of disease severity or a clinical diagnosis of AD [22]. However, since all DNA modifications at a locus can affect transcription, we hypothesized that conducting a more integrative analysis for all possible CpG modifications, as performed here, can be more informative in identifying disease-associated dysregulated loci. Notably, this approach highlighted two additional Braak stage-associated regions in *TNXB* and *DNAJB13* that were only identifiable by our DMoR analysis, and not when conducting any of the classical regional analysis as performed before.

By doing so, a gene-coding region in *TNXB* showed up as our most significantly dysregulated hit associated with Braak stage. Dysregulation was found both at the level of 5-uC and 5-mC, where an overall decrease in DNA methylation was related to increasing neuropathological features. The second region in *TNXB* that was identifiable only by our DMoR analysis, and which was located more downstream of the first Braak-associated region, displayed a similar increase in unmodified positions at all CpG sites, further emphasizing a potential role for active DNA demethylation in this gene in AD.

Interestingly, dysregulation at the level of DNA methylation in a similar region of *TNXB* has previously been observed in an AD EWAS targeting cortical tissues, including the entorhinal cortex, superior temporal gyrus and prefrontal cortex, which were derived from the exact same patient cohort as used in the present study [11]. *TNXB* expresses a glycoprotein with anti-adhesive properties, but its exact physiological role in the brain is not fully understood [27]. Next to epigenetic variation, genetic variation in this locus has also been directly associated with a risk of developing AD [28–30], as well as with progressive supranuclear palsy [31], which is a tau-related disorder.

In addition to AD, epigenetic dysregulation in a similar locus of *TNXB* has also been identified in EWAS targeting peripheral blood samples of patients and people at risk of developing several stress-related disorders, including anorexia nervosa, schizophrenia, remitted major depressive disorder and bipolar disorder [32–34]. These are interesting observations, given the established link between stress-related pathology and AD, especially when considering the physiological role of the DRN in the stress response. Evidently, more research on the exact function of TNXB, the interplay between genetic and epigenetic variation in its gene, and elucidating the exact cause-effect relationship underlying this AD neuropathology-associated epigenetic signature, is vital to develop a better understanding of its exact role in the disease.

Next to *TNXB*, another DMoR in the DRN bulk tissue that has previously been nominated in other AD EWAS was identified for *PGLYRP1*. The same gene displayed differential methylation in the aforementioned cross-cortex analysis using the same AD cohort as studied in the present manuscript [11]. Furthermore, a similar locus has been reported to demonstrate differential methylation in peripheral blood samples derived from individuals with down syndrome, a patient population known to be at increased risk of developing AD [35]. *PGLYRP1* encodes an innate immunity protein that is a known activator of TREM-1 [15]. Studies have demonstrated that overexpression of TREM-1 in APP/PSEN1 mice, a mouse model for AD, facilitates microglial-mediated amyloid beta clearance and restores AD-related cognitive impairment, emphasizing the importance of the PGLYRP1-TREM-1 interaction in the pathophysiology of AD [16]. Whether there is a functional connection between the reported epigenetic alterations in *PGLYRP1* and TREM-1 efficacy remains to be elucidated.

Besides *TNXB* and *PGLYRP1* that both have been identified in other AD EWAS, we also revealed novel regions in *RBMXL2*, *CAST*, *GNAT1*, *MALAT1* and *DNAJB13* displaying changes associated with AD neuropathology at the level of 5-uC and 5-mC. Interestingly, while representing new epigenetic associations, *CAST*, *MALAT1,* and *DNAJB13* have been functionally linked to AD before, supporting the relevance of our findings. Calpastatin, the protein expressed by *CAST*, is known to protect against neuronal death induced by amyloid beta [36, 37]. Furthermore, calpastatin depletion has been shown to act upstream of calpains to activate a calpain-dependent cascade of protein kinase activation, cytoskeletal protein hyperphosphorylation, cytoskeletal proeolysis and neurodegeneration [38]. In another study using APP/PSEN1 mice, calpastatin downregulation was linked to hyperglycemia and the promotion AD pathological hallmarks, the former of which is typically observed in diabetes, a known risk factor for AD [39]. The long noncoding RNA *MALAT1*, on the other hand, has recently been shown to convey neuroprotective effects in AD by inhibiting apoptosis and inflammation while promoting neurite outgrowth [40, 41]. Lastly, DNAJB13 belongs to the largest chaperone family regulating HSP40 chaperones, which have been reported to impede the amyloidogenic cascade by preventing amyloid beta aggregation [42, 43]. Moreover, the exact same region identified in our EWAS has previously been shown to be differentially methylated in peripheral blood samples of people with Down syndrome, further pointing towards a potential association with AD pathology [35]. Further research is necessary to assess whether the identified epigenetic alterations in these loci also have functional implications and, hence, could mediate the effects observed in the aforementioned studies. For *GNAT1* and *RBMXL*, no direct or indirect link or functional connection to AD has been identified to date.

In the GO term pathway analyses, we were furthermore able to identify altered biological mechanisms in the DRN bulk tissue related to our top-ranked DHPs. Strikingly, genes related to ‘homophilic cell adhesion via plasma membrane adhesion molecules’ and ‘calcium ion binding’ were overrepresented in this brainstem region. Interestingly, the cell adhesion molecules annotated to this aforementioned GO-term have a major function in dendrite development, synaptic connectivity and neural circuit formation [44, 45], closely linked to key neuropathological features of AD. In the context of ‘calcium ion binding’, an increasing numbers of studies suggests that disruption of intracellular calcium ion homeostasis plays important roles in orchestrating the dynamics of AD neuropathology and associated memory loss, as well as cognitive dysfunction [46]. Calcium dysregulation may even play an important role in the pathogenesis of AD, by inducing synaptic deficits, and by promoting the accumulation of amyloid beta plaques and neurofibrillary tangles [47]. All in all, these findings confirmed that the identified differentially modified genes are strongly linked to well-known AD-associated neuropathological processes, which further emphasizes a key role for epigenetic dysregulation in the DRN of AD patients.

Given that bulk tissue was used in the DRN EWAS, it is expected that various cellular populations residing in this brainstem region, including serotonergic neurons and other parenchymal cells such as glia, contribute to the obtained epigenetic signals. As such, the cellular heterogeneity of these bulk tissue samples constitutes a major source of noise in epigenetic profiling approaches, representing a reoccurring challenge in AD EWAS [22]. Furthermore, since AD is characterized by neuronal loss and increased microglia activation, differences in cellular compositions between tissue samples obtained at varying stages of the disease can interfere with the correct interpretation of the acquired epigenetic data. To tackle this issue and based on the hypothesized role of the serotonergic neurotransmitter system in mediating affective symptoms linked to *e.g.* early stages of AD, making use of a second independent cohort, we therefore isolated single serotonergic and non-serotonergic cells from the DRN by means of LCM, using an immunohistochemical labeling of 5-HT. Bisulfite methylation profiles in the previously Braak stage-associated region of *TNXB* were then quantified in both cell types obtained from healthy individuals and AD patients.

By doing so, we were able to demonstrate that the epigenetic signature in *TNXB* within this brainstem nucleus is strongly dependent upon the cell type analyzed. Whereas an AD-associated increase in DNA bisulfite methylation was found within 5-HT+ cells, a decrease in DNA bisulfite methylation was identified in patient-derived 5-HT-cells. As such, these data suggest that AD-associated epigenetic dysregulation in *TNXB* within the DRN is likely attributable to non-serotonergic cells, as the identified patterns in these cells overlapped with the previously obtained bulk tissue EWAS data. For *TNXB*, this notion is furthermore supported by another study [11] that has found dysregulation in this gene in several brain regions that are known to be free of serotonergic cell bodies. These cell-specific findings also illustrate that a potential loss of serotonergic neurons, as commonly observed in AD [2, 48], possibly resulting in a different proportion of various cell types within the bulk tissues examined, is not able to explain our bulk tissue EWAS data, indicating a true epigenetic AD-specific signature. Thus, our results support the notion that epigenetic data derived from heterogeneous post-mortem bulk tissue should be interpreted with caution, as changes in one cell type could negate or mask changes in (an)other(s).

### Concluding remarks

Taken together, the present study strongly implicates a role for *TNXB* dysregulation and other loci including *PGLYRP1*, *RBMXL2*, *CAST*, *GNAT1*, *MALAT1*, and *DNAJB13*, in the DRN of AD in the development and course of the disease. Moreover, for the first time, we highlighted potential cell-specific effects regarding *TNXB* that likely will have crucial implications for future planned epigenetic studies in AD. Our data warrants the need for single cell(-type) neuroepigenetic analyses, opposite to the more common bulk tissue analyses that have been performed to date in AD EWAS. Evidently, interrogation of epigenetic marks is most informative when studied at a single-cell level, where intercellular differences can be dissected leading to a more refined understanding of their contribution to the disease. We therefore strongly believe that future endeavors implementing single cell(-type) approaches combined with similar or more sophisticated epigenome-wide techniques as used here, will be increasingly valuable to dissect the exact contributions of epigenetic dysregulation in the etiopathogenesis of AD.

## Materials and methods

### Study sample description

The post-mortem DRN bulk tissue used in the discovery EWAS and technical validation analysis using bisulfite pyrosequencing was obtained from the Medical Research Council (MRC) London Brain Bank for Neurodegenerative Disease (London, UK). Samples were provided with informed consent according to the Declaration of Helsinki (1991) and ethical approval for the study was provided by the NHS South East London REC 3. For the EWAS, 93 DRN samples from both AD patients and neurologically healthy controls were used. Upon screening for AD neurofibrillary tangles at autopsy, individuals were selected based on Braak stage ranging from 0 to VI. All samples were dissected by trained specialists, snap-frozen, and stored at −80°C up until further processing.

Post-mortem DRN samples used in the cell-specific validation analysis were obtained from a second and independent cohort derived from the BSHRI (Sun City, Arizona, USA). Age-matched samples from 12 females without-(Control cases) and 12 females with a clinical diagnosis of AD (AD cases) were collected. Each individual and her respective relative(s) consented to brain autopsy for scientific research as part of the BSHRI BBDP [24]. Written informed consent for the autopsy was obtained in compliance with institutional guidelines. The institutional review board approved the entire study, including the recruitment, enrollment, and autopsy procedures. A final diagnosis of AD at autopsy was made by following the National Institutes of Health (NIH) AD Center criteria [24]. Comorbidity with any other type of dementia, mild cognitive impairment, cerebrovascular disorders, and the presence of non-microscopic infarcts were applied as exclusion criteria. All samples were dissected by trained specialists, snap-frozen and stored at −80°C up until further processing. Demographic and relevant information about the sample cohorts are provided in Table 1.

### Genome-wide DNA modification profiling

Genomic DNA was extracted from ∼50 mg of DRN tissue using a standard phenol-chloroform extraction method. The TrueMethyl 24 Kit version 2.0 by CEGX (Cambridge Epigenetix Limited, Cambridge, UK) was used for bisulfite- and oxidative bisulfite treatment of the DNA isolated from DRN bulk tissue. All laboratory procedures were performed at GenomeScan (GenomeScan B.V., Leiden, the Netherlands), according to the manufacturer’s instructions. Before bisulfite treatment, high molecular weight DNA was quantified using a PicoGreen assay (Invitrogen, Carlsbad, California, USA) and the DNA quality was assessed by gel-electrophoresis, confirming that all samples were of sufficient quantity and quality. From each individual, 1 µg of isolated DNA was used, which, after purification and denaturation, was split up into two samples that underwent either the DNA oxidation or a mock DNA oxidation treatment, for oxidative bisulfite- and customary bisulfite-treated samples, respectively. Subsequently, all samples were bisulfite-converted and the yield of the (oxidative) bisulfite DNA was assessed by a Qubit ssDNA assay (Invitrogen). An additional restriction quality control was performed for qualitative assessment of 5-hmC oxidation and bisulfite conversion using the Fragment Analyzer. From each bisulfite- and oxidative bisulfite-treated DNA sample, 8 µL was amplified and hybridized on the Infinium MehtylationEPIC BeadChip (EPIC array; Illumina, Inc., San Diego, CA, U.S.A.). All samples were randomized in a balanced manner for sex and Braak stage, and bisulfite- and oxidative bisulfite-treated samples from the same individuals were run on the same chip to avoid batch effects. The Illumina iScan was used for imaging the arrays. Sample preparation, hybridization, and washing steps were performed according to the manufacturer’s instructions.

Raw signal intensities generated from both bisulfite- and oxidative bisulfite-treated samples were used to construct *MethylumiSet* objects using the ‘*readEPIC*’ function in the *wateRmelon* package [49] and *RGChannelSet* objects using the *‘read.metharray.exp*’ function in the *minfi package* [50]. By using 59 single nucleotide polymorphism (SNP) probes on the array, we were able to confirm that the matching samples that underwent bisulfite and oxidative bisulfite treatment were sourced from the same DNA samples. Probes with common (minor allele frequency (MAF) > 5%) SNPs in CG or single base extension positions or cross-reactive probes were flagged and discarded. A principal component analysis (PCA)-based approach was used to examine a potential mismatch between reported and predicted sex. By using the ‘*pfilter’* function within the *wateRmelon* package, samples with a detection *p* >0.05 in more than 5% of probes, probes with more than three beadcounts in 5% of the samples, and probes having 1% of samples with a detection *p* value >0.05 were identified and removed. Next, the bisulfite and oxidative bisulfite datasets were split, and by using the ‘*preprocessRaw*’ function, the red and green channels for the Illumina methylation arrays were converted into methylation signals followed by a *Noob* background correction method with dye-bias normalization [51]. In order to estimate the proportion of DNA modifications, the maximum likelihood (ML) method described by Qu *et al.* [23] was used. The MLML function within the *MLML2R* package [52] was applied, which used the combined methylated and unmethylated signals from the bisulfite and the oxidative bisulfite arrays as an input, and then returned the estimated proportion of 5-uC, 5-mC, and 5-hmC for each CpG site. Only probes with a mean beta value >0.1 on both the bisulfite- and oxidative bisulfite EPIC arrays were included in the MLML method. Finally, for 5-hmC, only sites where 5-hmC was present in more than half of the sample population were used for the analysis.

### Bisulfite pyrosequencing

For the technical validation analysis, 200 ng of DNA was used from a subset of 62 isolated DNA samples that were previously utilized for the EPIC arrays. The EZ-96 DNA Methylation-Gold Kit (D5008; Irvine, CA, USA) was used for bisulfite conversion. Finally, bisulfite-converted DNA samples were eluted in 20 µl of elution buffer, and 1 µl was used for polymerase chain reaction (PCR) amplification with subsequent bisulfite pyrosequencing. Bisulfite methylation profiles, *i.e.* 5-mC+5-hmC, levels were quantified across eight individual CpG sites in *TNXB*, including cg10365886, cg14188106, and cg07524919, spanning from 32063869 to 32063940 on chromosome (chr) 6 (Ensembl GRCh37 assembly). A single amplicon of 209 bp (chr6:32063788-32063996) was PCR-amplified using primers designed with the PyroMark Assay Design software 2.0 (Qiagen, Hilden, Germany) (Supplementary Table 10). PCRs were performed with an initial denaturation step at 95°C for 5 min, followed by 45 cycles at 95°C for 30 sec, 62°C for 30 sec, and 72°C for 30 sec, with a final extension step at 72°C for 1 min. All PCR reactions, contained 2.5 μl PCR buffer (10X) with 20 mM MgCl2, 0.5 μl 10 mM dNTP mix, 1 μl of each primer (5 μM stock), and 0.2 μl (5 U/μl) FastStart™ Taq DNA Polymerase (Roche Diagnostics GmbH, Mannheim, Germany) in a total volume of 25 μl. Pyrosequencing of the amplicon was performed using a single sequencing primer covering the 72-bp region. DNA bisulfite methylation was quantified using the PyroMark Q48 Autoprep system (Qiagen) and the Pyro Q48 Autoprep 2.4.2 software following the manufacturer’s instructions. Adequate sensitivity of the assay was initially confirmed using methylated and unmethylated DNA standards from the EpiTect PCR Control DNA Set (Qiagen).

### Laser capture microdissection

Laser capture microdissection was performed as described previously [25] to isolate serotonin-positive (5-HT+) and 5-HT-negative (5-HT-) cells from the DRN. In brief, frozen tissue sections of 10 μm were mounted onto polyethylene naphthalate (PEN) slides and fixed in ice-cold 50% acetone/50% ethanol solution for 5 min on ice. Sections were washed in ice-cold phosphate-buffered saline (PBS), blocked in 1% hydrogen peroxide for 2 min, followed by 3 quick submersions in ice-cold PBS. Sections were then incubated with a primary antibody against serotonin (5-HT; Abcam, ab66047; in PBS, 1:10) for 10 min at room temperature. After incubation, sections were washed three times with PBS and incubated with an avidin-biotin complex in PBS for 10 min at room temperature. Next, sections were washed three times in 50 mM Tris buffer and immersed in a 3.3’-diaminobenzidine (DAB) solution (4 ml 50 mM Tris; 100 μl DAB (10 mg/ml); 400 μl saturated nickel; and 4 μl of 5% hydrogen peroxidase) for 5 min, followed by two quick rinses in 50mM Tris to stop the reaction. All sections were stored at −80°C until further processing.

Since 5-HT is a monoamine neurotransmitter that is specifically expressed by serotonergic neurons [53], LCM of 5-HT-positive (5-HT+) cells from the DRN sections was performed based on 5-HT-immunoreactivity. Sections were dipped in 100% ethanol, allowed to dry, and loaded onto a Leica AS-LMD LCM microscope (Leica, Wetzlar, Germany). Single 5-HT+ cells were cut and then dropped into an inverted microcentrifuge cap containing 10 μl of Tris-EDTA (TE) buffer. Per individual, 150 5-HT+ cells were captured at 20X magnification and divided into three pools of 50 cells per microcentrifuge tube per person. For the non-serotonergic cells, *i.e.* 5-HT negative (5-HT-) cells, a similar amount of tissue devoid of 5-HT-immunoreactivity was collected from the exact same samples (Control, n = 10; AD, n = 10). All tissue isolated by LCM was stored at −80°C until further processing.

### Limiting dilution bisulfite pyrosequencing

DNA from each pool of 5-HT+ or 5-HT-cells was isolated and bisulfite-converted using the EZ DNA Methylation-Direct Kit (Zymo Research, Irvine, CA, USA) with the following adaptations, and as described previously [25]. In brief, 1 µl of proteinase K (20 µg/µl) and 11 µl of M-Digestion buffer (2X) were added to a microcentrifuge tube containing the cells and incubated overnight at 50°C. Subsequently, the complete lysate was transferred to a PCR tube and 143 µl of bisulfite conversion reagent was used to wash out the digestion tube before adding it to the sample. Bisulfite conversion was performed in a thermal cycler running at 98°C for 8 min and then at 64°C for 3.5 h. A volume of 200 µl binding buffer was added to the spin column before loading the bisulfite-converted sample. The PCR tube used for the bisulfite conversion was washed out twice by first adding 200 µl of binding buffer to the tube and then by transferring this volume to the sample-containing column. After centrifugation (10,000 x g; 30 sec), the column was washed with 100 µl washing buffer, incubated for 15 min with 200 µl desulfonation buffer, and washed twice again with 200 µl washing buffer. The bisulfite-converted DNA was eluted in a single Eppendorf tube by running 20 µl of elution buffer through the column twice (Two times at 10,000 x g; 30 sec). Eppendorf LoBind microcentrifuge tubes (Merck KGaA, Darmstadt, Germany) and TipOne Low Retention Tips (STARLAB, Hamburg, Germany) with low affinity for DNA were used throughout the whole procedure. PCR amplifications were performed directly after elution of the bisulfite-converted DNA.

The bisulfite pyrosequencing assays for *TNXB* used in the cell-specific validation analysis were based on a semi-nested PCR. Primers were designed with the PyroMark Assay Design 2.0 software (Qiagen), based on the Ensembl GRCh37 assembly (See Supplementary Table 10). Only for the LDBSP analysis [25], bisulfite-treated DNA from a pool of 5-HT+ cells was diluted to a single allele level by adding a PCR mixture with a capacity of 22 individual reactions to the sample (determined empirically). Each PCR reaction made use of 2.5 μl PCR buffer (10X) with 20 mM MgCl2, 0.5 μl 10 mM dNTP mix, 1 μl of each primer (10 μM stock), and 0.2 μl (5 U/μl) FastStar Taq DNA Polymerase (Roche Diagnostics GmbH, Mannheim, Germany) in a total volume of 25 μl. After adding the bisulfite DNA to the complete mixture, the sample was pipetted up and down in order to homogeneously disperse all bisulfite-converted DNA molecules throughout the solution, and fractions of 25 µl were divided over 22 wells of a microtiter plate. An amplicon of 440 bp (chr6:32063558-32063997), using the outer primers for *TNXB*, was amplified based on an initial denaturation step at 95°C for 5 min, followed by 43 cycles at 95°C for 30 sec, 56°C for 30 sec, and 72°C for 1 min, with a final extension step at 72°C for 7 min. For the nested PCR reaction, 1 µl of the multiplex product was used as a template. An amplicon of 432 bp (chr6:32063566-32063997) for *TNXB* was then amplified using the inner primers with the same PCR compound concentrations and under similar thermocycler conditions as described above, although with 45 cycles and an annealing temperature of 58°C. Bisulfite-treated DNA derived from pools of 5-HT-cells was also amplified using a semi-nested PCR, *i.e.* by extended pre-amplification, although with omitting the step of over-diluting the bisulfite-converted DNA pool to single alleles, since the exact cell numbers could not be determined from the same tissue devoid of 5-HT-immunoreactivity. All PCR settings, as well as subsequent bisulfite pyrosequencing procedures, were performed under the same conditions for both of the analyzed cell types.

Bisulfite pyrosequencing was used to determine the bisulfite methylation, *i.e.* cumulative signal of 5-mC and 5-hmC, status of the first Braak stage-associated region in *TNXB* in both 5-HT+ and 5-HT-cells of the DRN. The DNA bisulfite methylation status of *TNXB* was quantified over a region spanning from 32063774 to 32063913 within chromosome 6 (Ensembl GRCh37 assembly), including six Illumina probes: cg10365886, cg07524919, cg14188106, cg26266427, cg01337207, and cg02989255. To maximize CpG site coverage over the targeted regions, three pyrosequencing primers were used (See Supplementary Table 10). The PyroMark Q96 MD pyrosequencing system (Qiagen) with the PyroMark Gold Q96 CDT reagent kit (Qiagen) was used according to the manufacturer’s instructions. Bisulfite methylation levels at a single CpG site resolution were quantified with the Pyro Q-CpG 1.0.9 software (Qiagen). Adequate sensitivity of the assays was assessed as described earlier using (un)methylated DNA standards from the EpiTect PCR Control DNA Set (Qiagen).

LDBSP data processing was performed as described previously [25]. In brief, sequenceable amplicon-yielding PCR reactions for *TNXB* were assessed for their representation of single-, two or three-alleles. Assessment was performed using thresholds of (1) ≤8.33% and ≥91.33%, (2) 50±8.33%, and (3) 33.33±8.33% and 66.66±8.33%, and based on a subsequent CpG-site-dependent calling procedure, combined with an integrated in-depth analysis of the raw CpG methylation rates. All PCR products were thoroughly inspected by two investigators that were blinded to the experimental conditions, and a decision on the total number of alleles present in each individual reaction, *i.e.* one, two or three alleles, was made independently, while taking into account the criteria described in [25]. A cross comparison between the independent *TNXB* score sheets was then performed (98.73% overlap in scoring) and reactions with a deviating score between the first two investigators were assessed by a third (blinded) investigator. A final allele number was then assigned for these reactions based on the overlap between the score sheets of the third and first two investigators, *i.e.* when two out of the three investigators assigned the same score, then this allele number was used for the respective reaction. For each individual, the methylation rate for each CpG site was determined by expressing the number of methylated CpG sites as a percentage of the total number of CpG sites over the estimated alleles, whilst correcting for the number of alleles present in each of the reactions, *i.e.* one, two or three alleles [25]. Further LDBSP parameters can be found in Supplementary Table 11.

### Statistical analysis

All computations and statistical analyses were performed using R version 3.3.2 [31]. In the DRN discovery EWAS, DMoPs associated with Braak stage at the level of unmodified (5-uC) methylated (5-mC), and hydroxymethylated (5-hmC) cytosine positions, i.e., DUPs, DMPs and DHPs, were identified using multiple linear regression models for each individual CpG probe. Surrogate variable analysis (SVA) was performed to determine and estimate variation introduced by cell type heterogeneity in the bulk tissue. First five SVs along with age and sex were included as covariates in the models. The inflation index lambda (λ) was computed for the p values obtained from each EWAS and QQ plots. The *p* values were adjusted for multiple testing using the Benjamin-Hochberg FDR procedure. Illumina EPIC array probes were annotated using the Illumina UCSC gene annotation (Ensembl GRCh37 assembly).

We used the comb-p tool [32] to perform regional analysis based on spatial correlation of the *p* values determined by each EWAS. A sliding window of 500bp and a seeding p value of 1.0E-3 were set for the regional analyses and the Šidák correction was used for multiple testing correction. Additionally, we employed the comb-*p* algorithm to identify DMoRs. For this purpose, we utilized the smallest *p*-values obtained from either 5mC, 5hmC, or 5UC as input to the comb-*p* algorithm. Consequently, DMoRs represented clusters of neighboring CpG sites exhibiting associations with any type of modification within a specific region, rather than clusters displaying associations exclusively within a particular type of modification (as observed in the case of DURs, DMRs, and DHRs).

For the top-ranked DUPs, DMPs and DHPs, underlying biological processes and pathways were examined using a Gene Ontology (GO) enrichment analysis. Analyses were performed using the *missMethyl* package [54], which accounts for the different number of probes annotated to a single gene on the array.

For the pyrosequencing validation analysis of *TNXB* in the DRN bulk tissue, CpG methylation data that passed quality control were exported from the Q48 Autoprep 2.4.2 software to the R statistical environment. Subsequently, a multiple linear regression analysis was performed with the bisulfite methylation signals per CpG site as outcome, Braak Stage as predictor, and with age and sex added as covariates in the model (similar to the EWAS described above). Cases with missing CpG bisulfite methylation values after pyrosequencing (n = 2) were excluded pairwise. In addition, we conducted Pearson correlation analyses to examine the correlation between bisulfite methylation values obtained from bisulfite pyrosequencing with their corresponding three Illumina probes in TNXB (cg10365886, cg14188106, and cg07524919). For these technical validation analyses, a *p* value <0.05 was considered statistically significant.

For the cell-specific validation analysis, average bisulfite methylation levels across the targeted region of *TNXB*, were calculated for both the 5-HT+ and the 5-HT-cells. For the 5-HT+ cells, the bisulfite methylation rates obtained for the targeted CpG sites were first averaged over the region per subject and subsequently over the experimental groups. For the 5-HT-cells, only cases for which all CpG sites surpassed quality control by the Pyro Q-CpG 1.0.9 software were included and averaged per subject, as well as the experimental groups. One AD case and one control case were excluded from the analysis (5-HT-cells; Control, n = 9; AD, n = 9), as these did not meet the quality control criteria. A general linear model with experimental condition (Control and AD cases) as between-subject factor and cell type (5-HT+ and 5-HT-cells) as within-subject factor was conducted to test for interaction and main effects. The data were analyzed using a general linear model based on experimental condition and cell type. A *p* value <0.05 was considered as statistically significant.

## Author Contributions

E.P., T.H., A.D.S, R.D., M.E., G.K., B.R., K.P.L, S.D.G, P.C., J.M., D.M., A.R., K.L, and D.L.A.v.d.H conceptualized the project. R.J.M.R., J.N., and A.I. conducted the experiments. R.J.M.R., E.P., A.I., J.R., R.L., and D.L.A.v.d.H. performed the data analyses. D.L.A.v.d.H., K.L., T. H., and D.M. provided the resources for the project. R.J.M.R., E.P, and D.L.A.v.d.H., wrote the manuscript, with critical revision by all other authors. D.L.A.v.d.H, and K.L. acquired the funding for the project. All the authors have read and agreed to the published version of the manuscript.

## Acknowledgements

This research was funded by by the Joint Programme—Neurodegenerative Disease Research (JPND) for the EPIAD consortium (http://www.neurodegenerationresearch.eu/wp-content/uploads/2015/10/Factsheet_EPI-AD.pdf) (DvdH). The project is supported through the following funding organizations under the aegis of JPND; The Netherlands, The Netherlands Organisation for Health Research and Development (ZonMw); United Kingdom, Medical Research Council; Germany, German Federal ministry of Education and Research (BMBF); Luxembourg, National Research Fund (FNR). This project has received funding from the European Union’s Horizon 2020 research and innovation programme under Grant Agreement No. 643417.

---

**Supplementary Figure 1.**
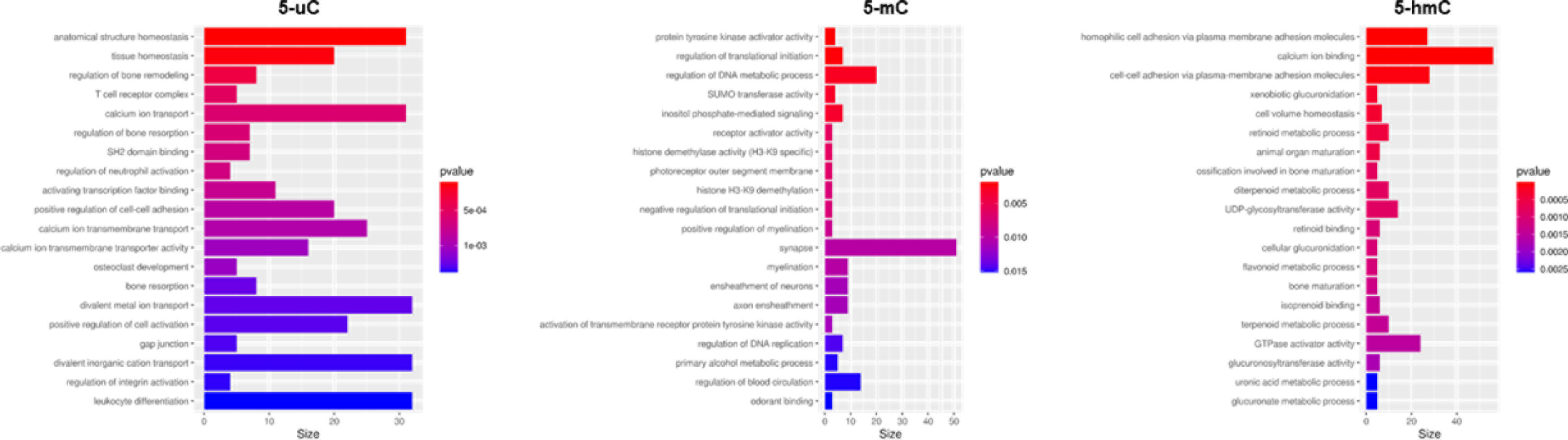
Gene Ontology (GO) terms enriched in the dorsal raphe nucleus (DRN). Displayed are the top 20 enriched GO terms in the DRN for each cytosine state: unmodified cytosine (5-uC), 5-methylcytosine (5-mC) and 5-hydroxymethylcytosine (5-hmC). The x-axis displays the number of differentially modified genes in the pathway.

**Supplementary Table 1.**
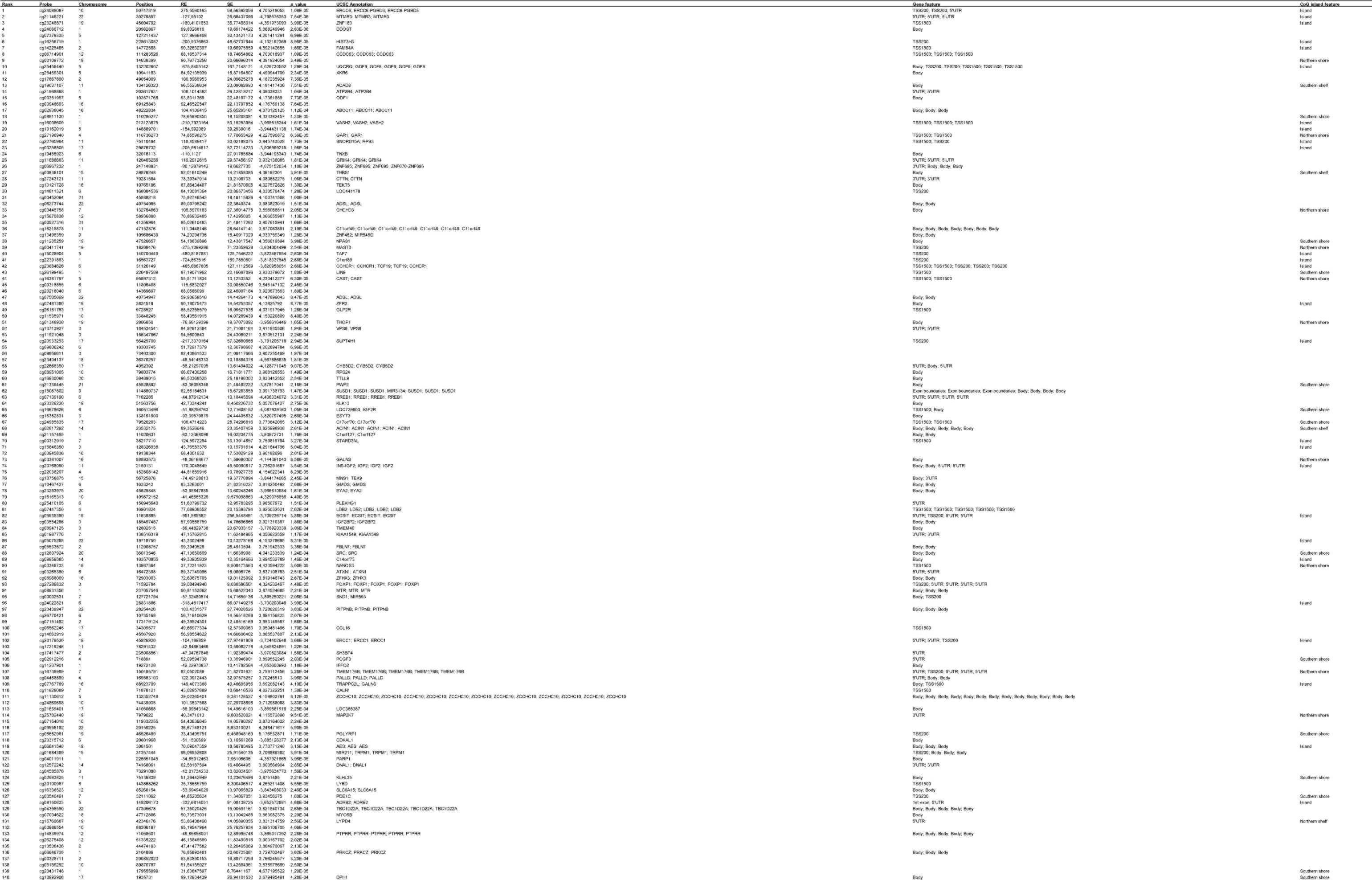
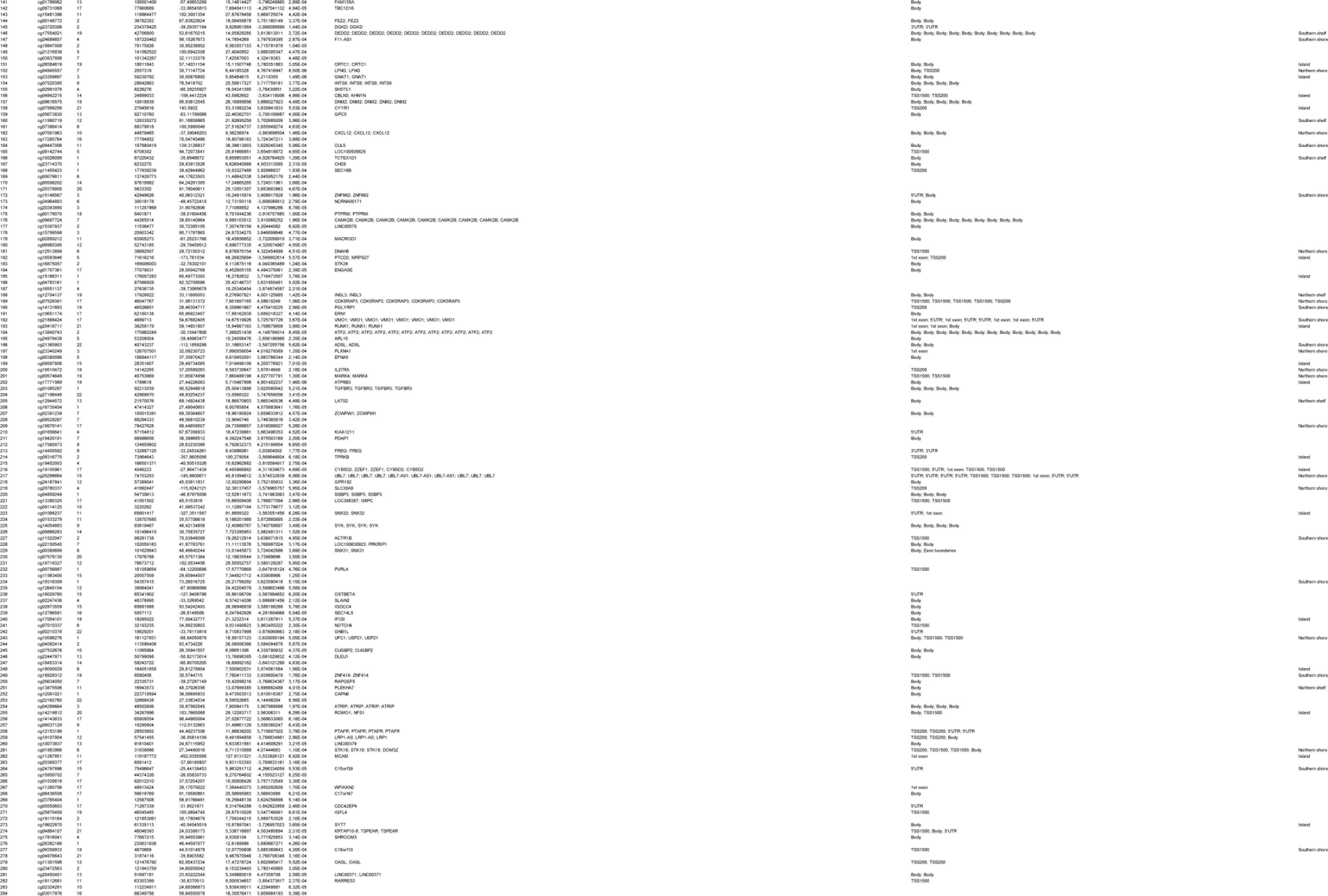
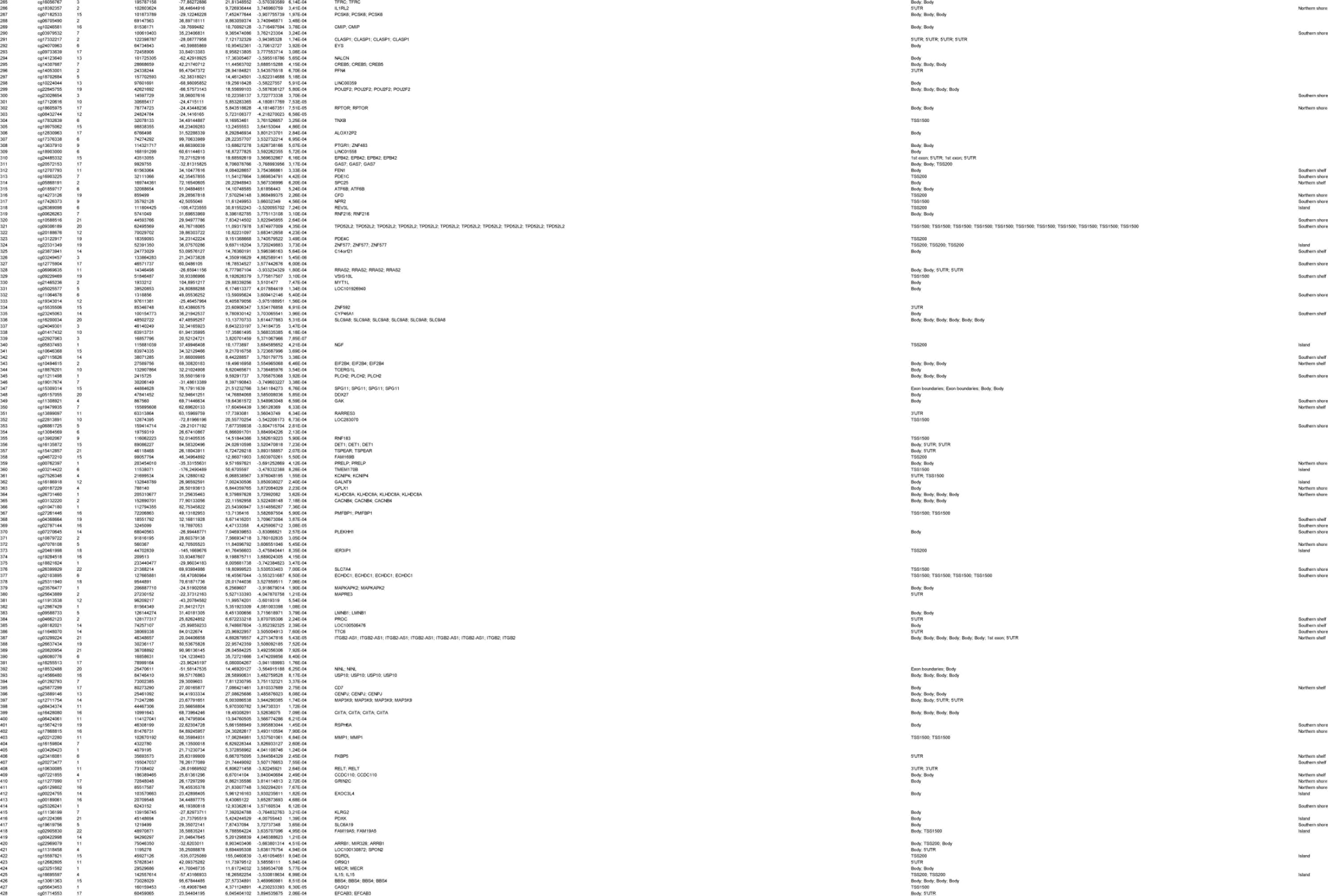
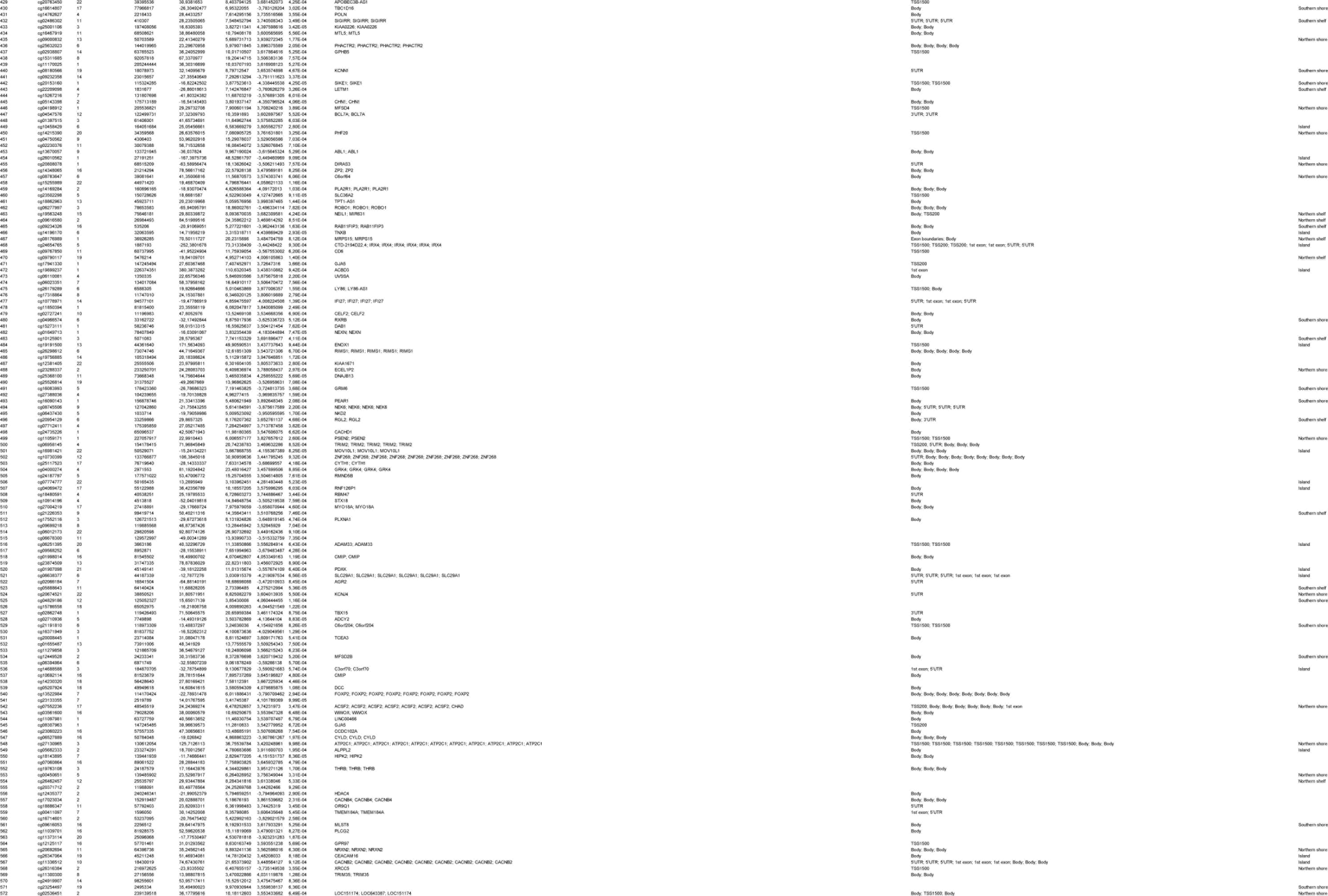
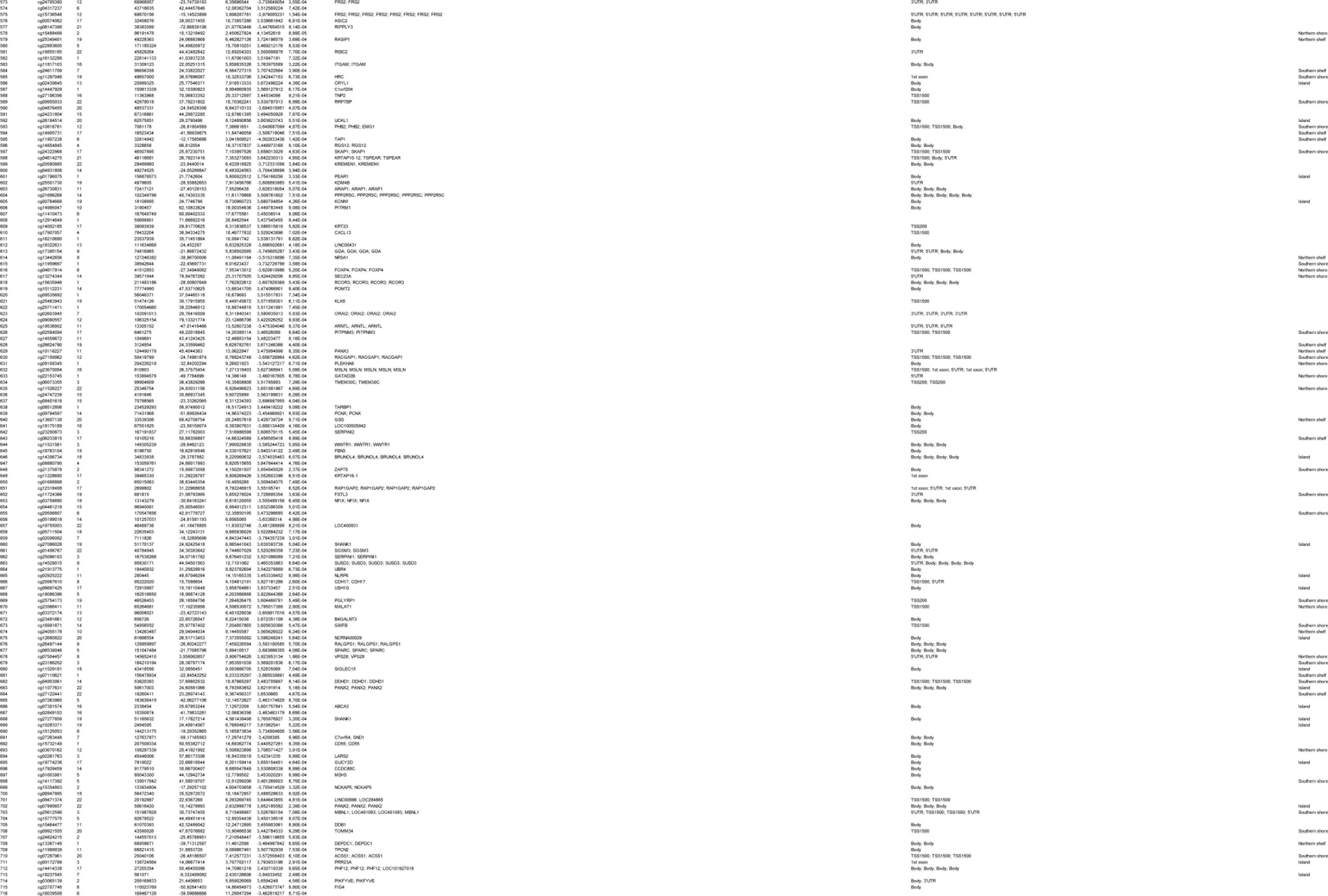
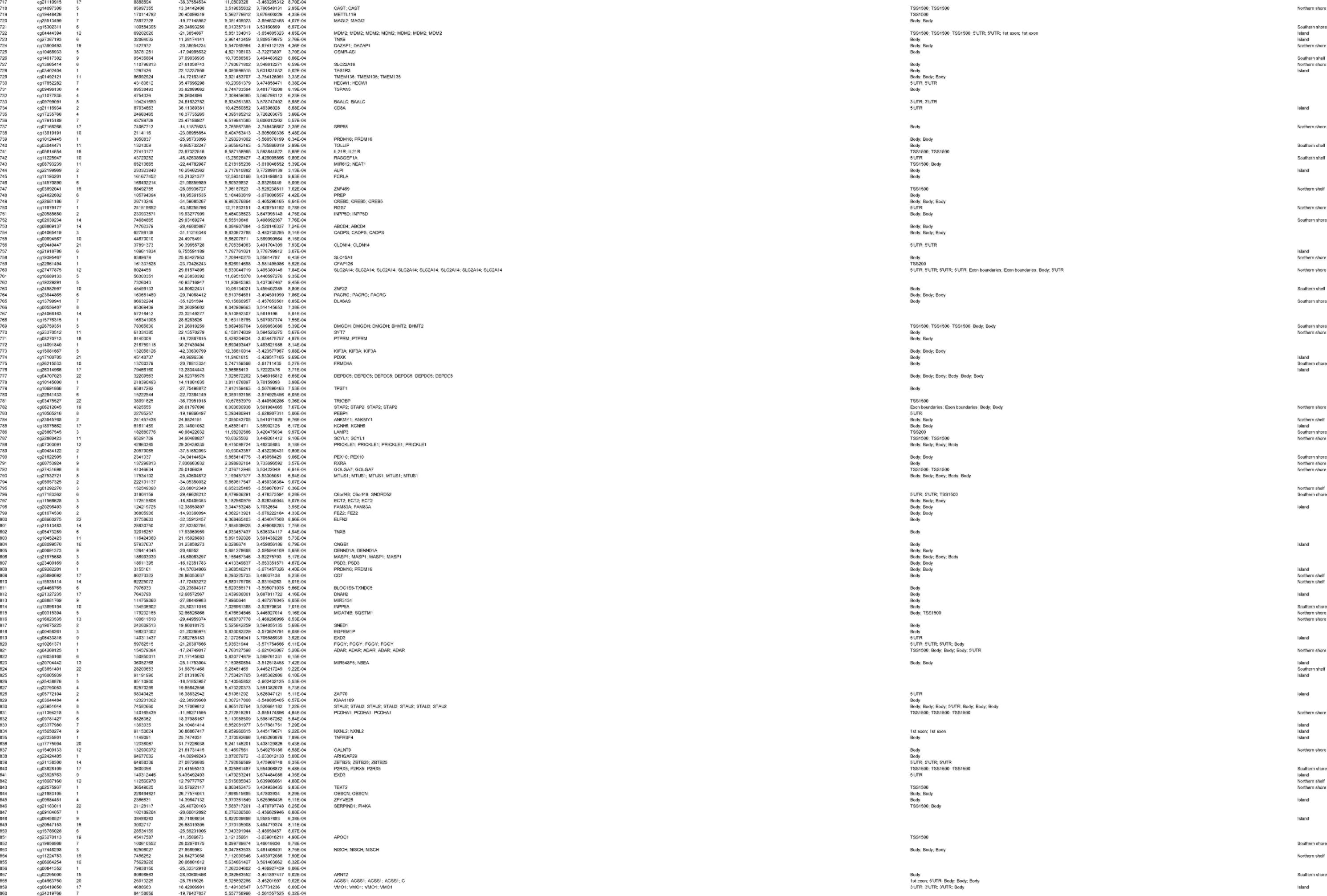
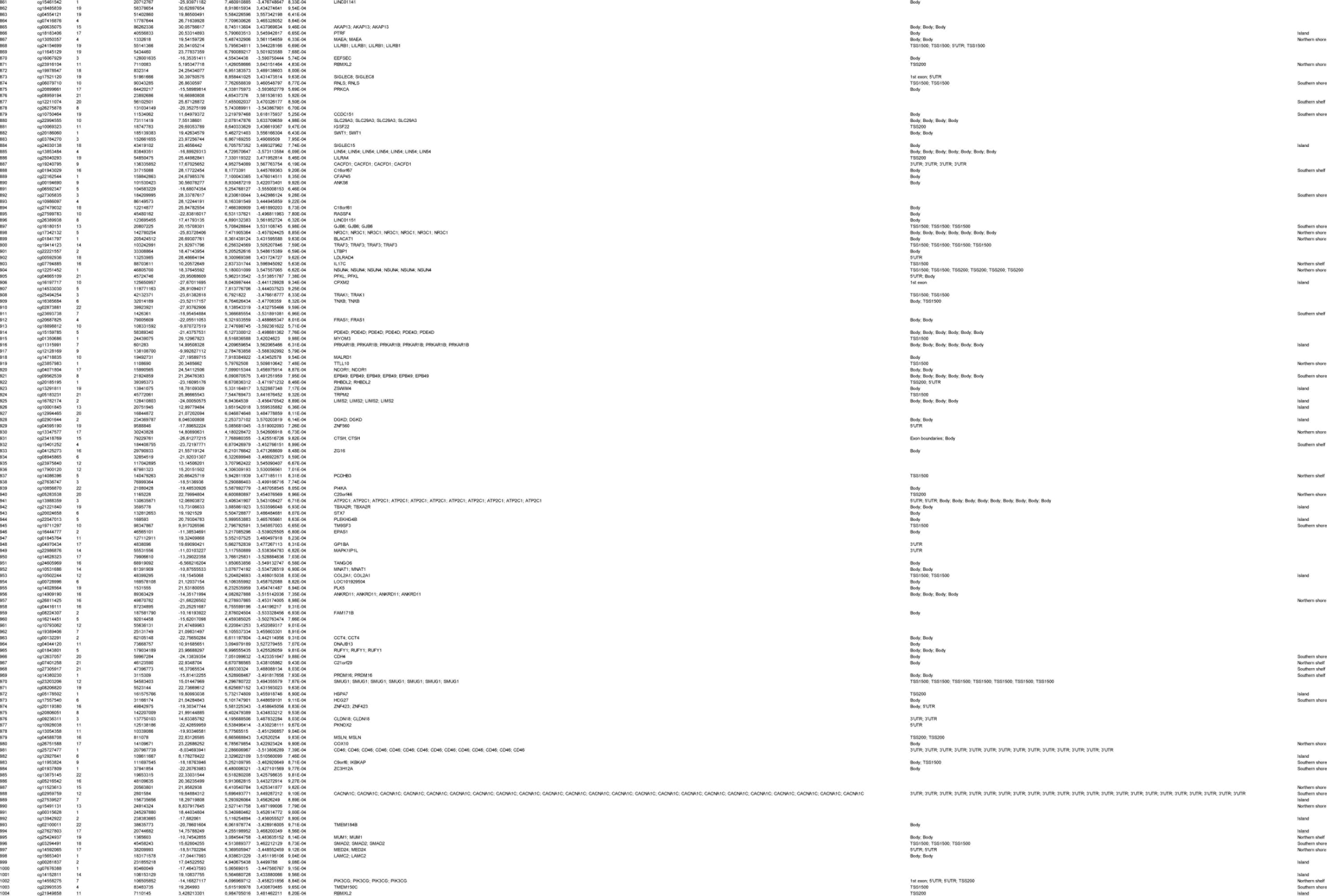
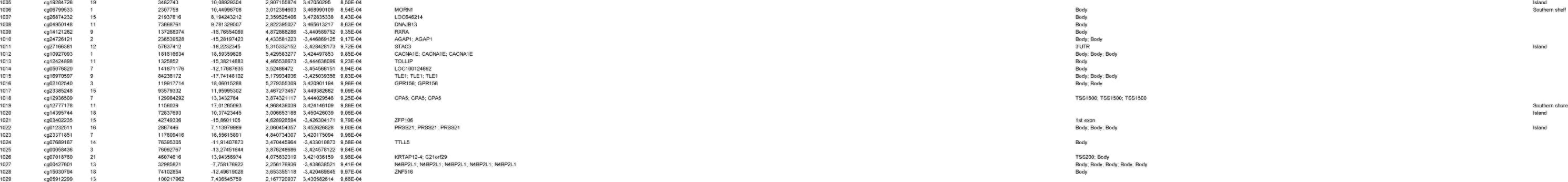
Top ranked differentially unmodified positions (DUPs) in the dorsal raphe nuclei (DRN) bulk tissue. Displayed for each ranked probe is the chromosomal location and position (Ensembl GRCh37 assembly), the regression estimate (RE) for the Braak stage-associated analysis, the standard error (SE), the t-statistics, the accompanying p values, the Illumina gene annotation (UCSC Annotation), the gene feature (TSS1500, 200 to 1500 nucleotides (nt) upstream of transcription start site (TSS); TSS200, up to 200 nt upstream of TSS; 5′UTR, 5′untranslated region; Body, gene body; 3′UTR, 3′ untranslated region) and the cytosine-phosphate-guanine (CpG) island feature. Probes are ranked based on a combined p value (cut-off = p < 0.001) and regression estimate ranking.

**Supplementary Table 2.**
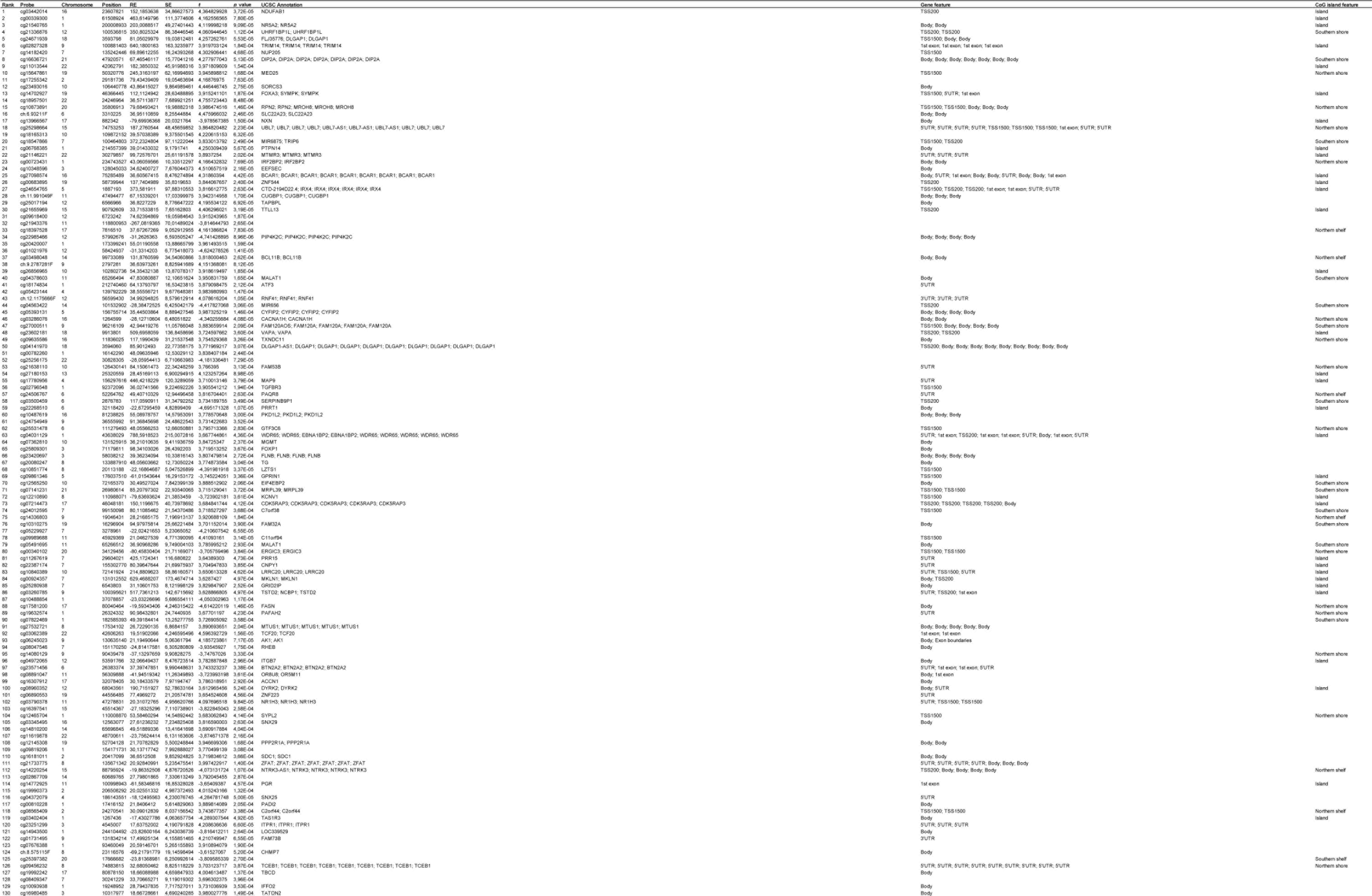
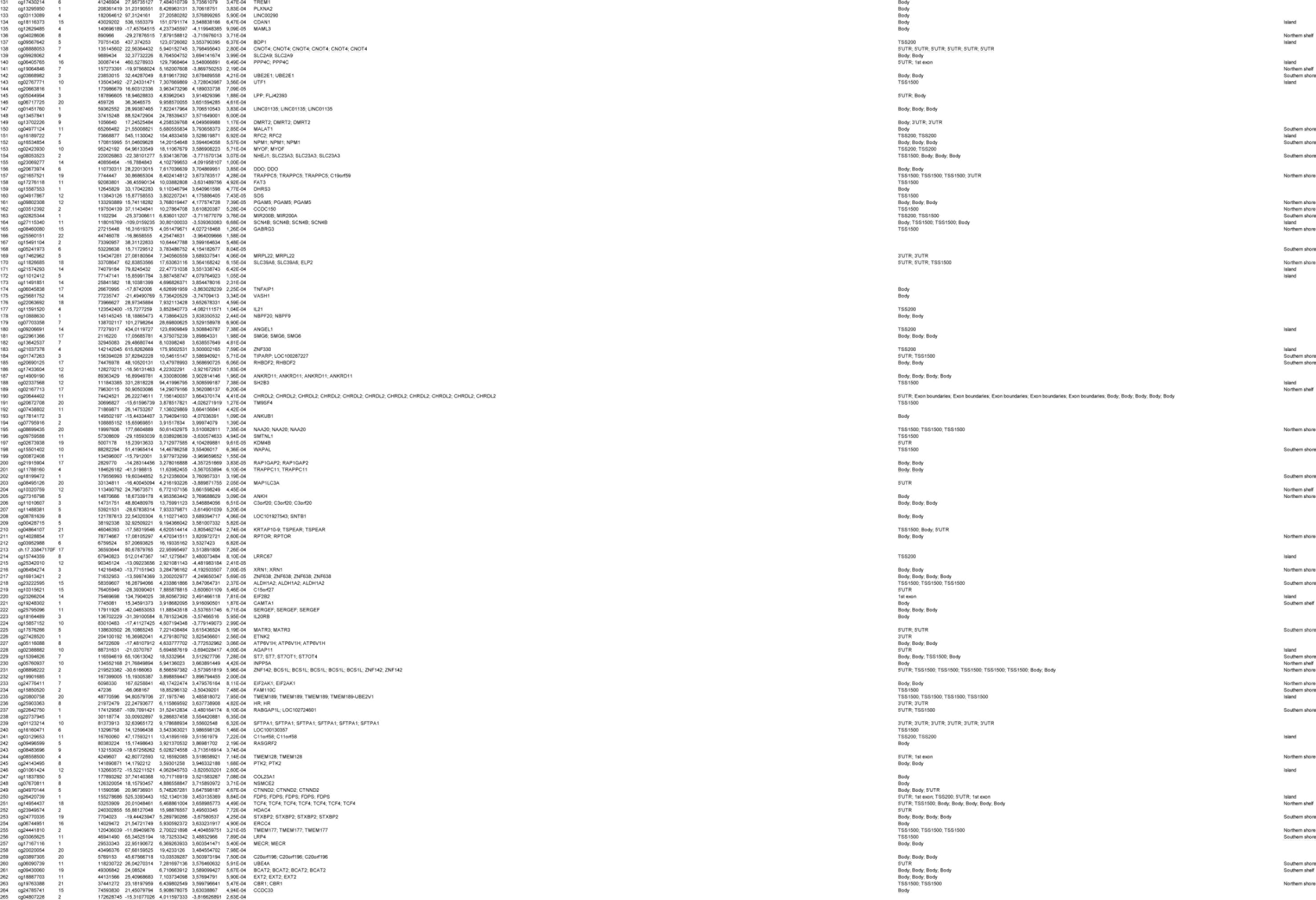
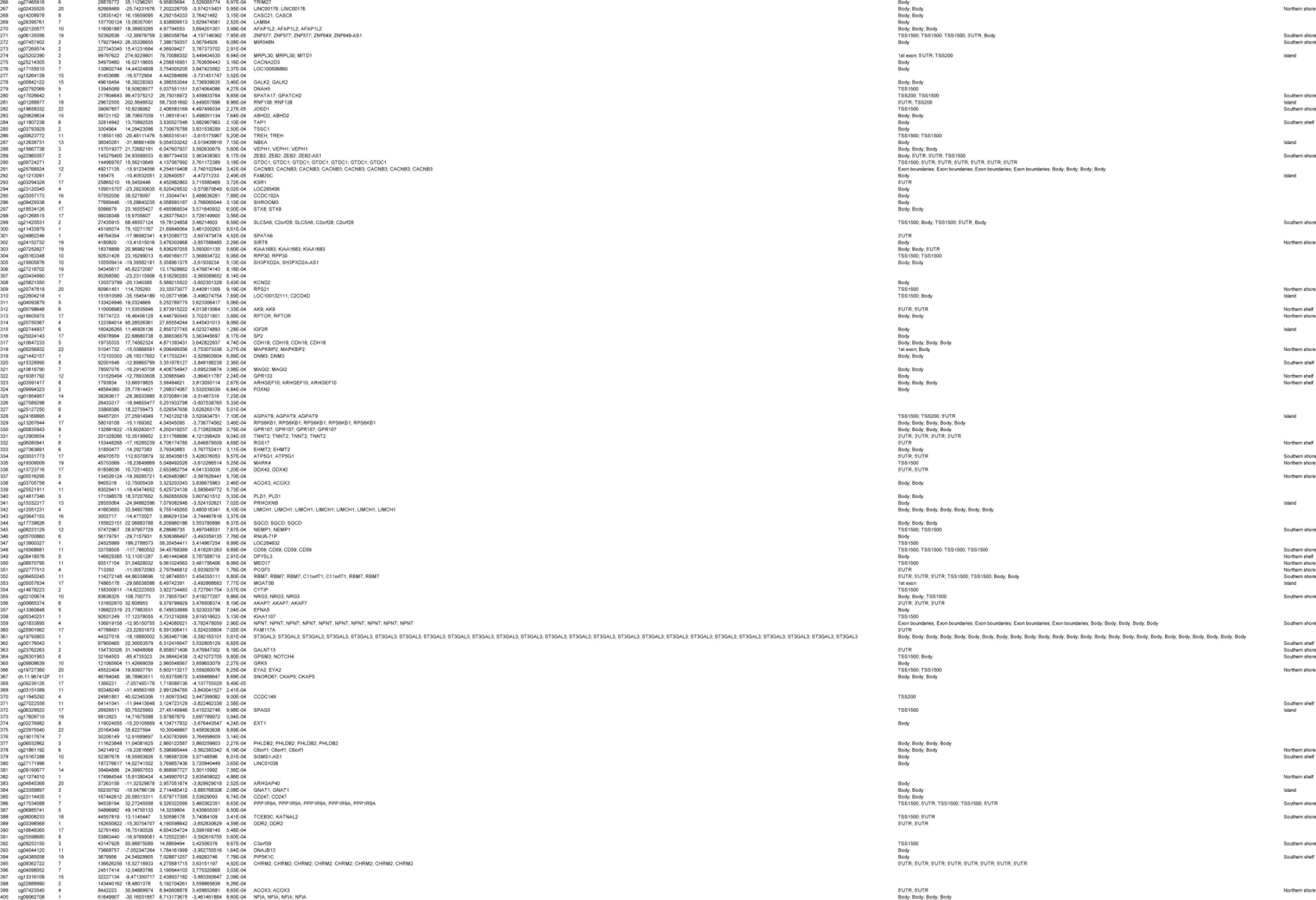
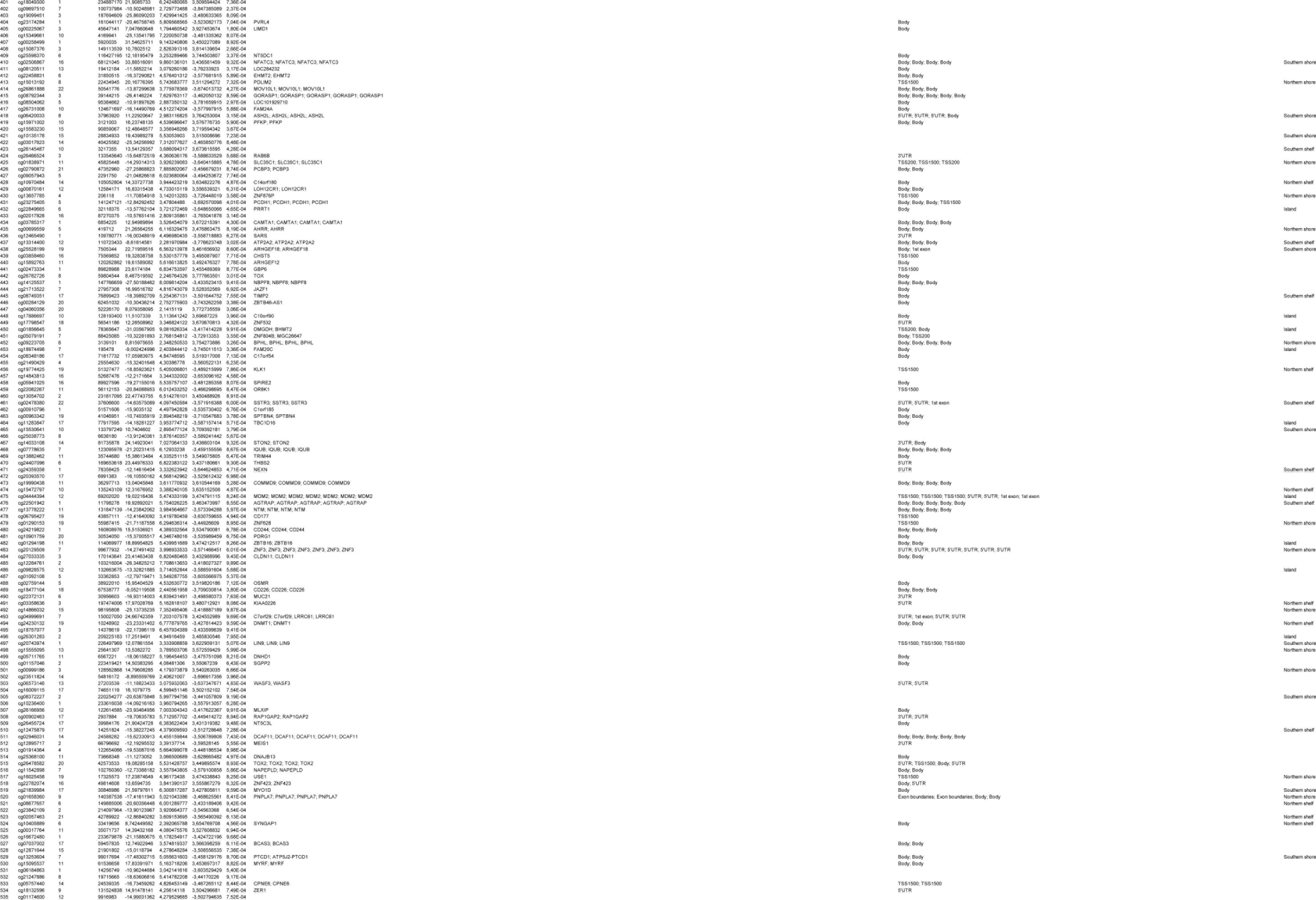
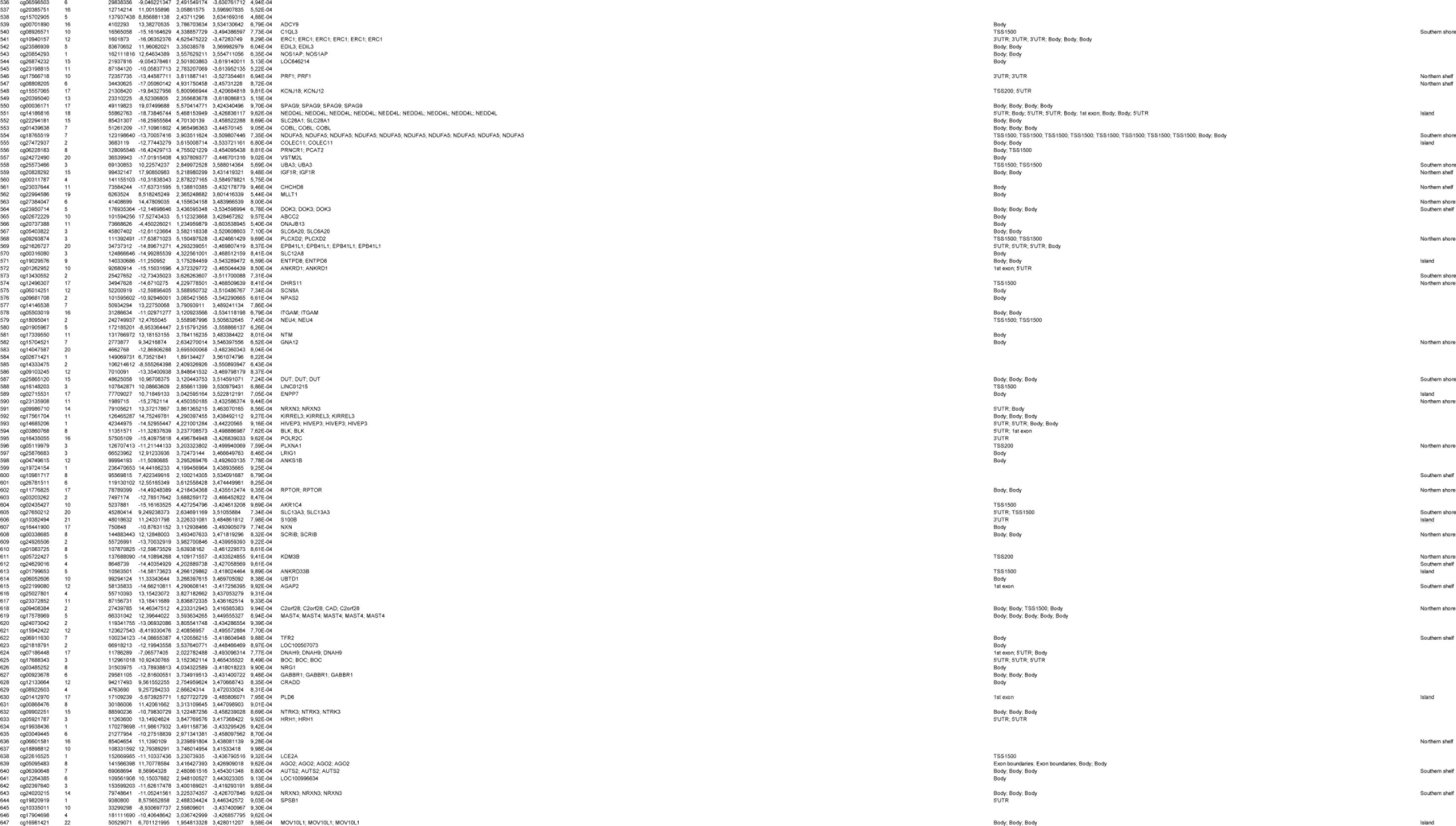
Top ranked differentially methylated positions (DMPs) in the dorsal raphe nuclei (DRN) Displayed for each ranked probe is the chromosomal location and position (Ensembl GRCh37 assembly), the regression estimate (RE) for the Braak stage-associated analysis, the standard error (SE), the t-statistics, the accompanying p values, the Illumina gene annotation (UCSC Annotation), the gene feature (TSS1500, 200 to 1500 nucleotides (nt) upstream of transcription start site (TSS); TSS200, up to 200 nt upstream of TSS; 5′UTR, 5′untranslated region; Body, gene body; 3′UTR, 3′ untranslated region) and the cytosine-phosphate-guanine (CpG) island feature. Probes are ranked based on a combined p value (cut-off = p <0.001) and regression estimate ranking.

**Supplementary Table 3.**
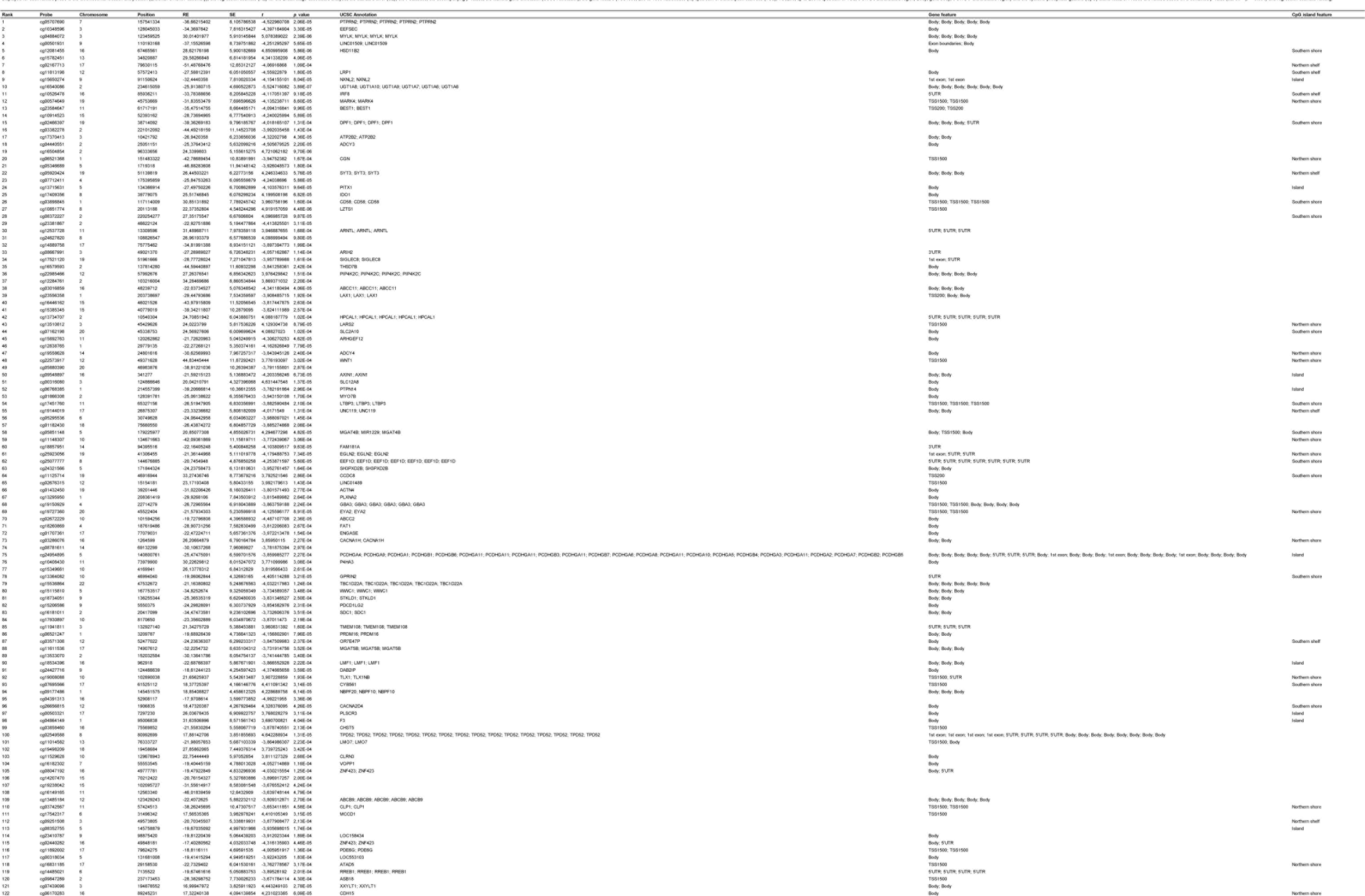
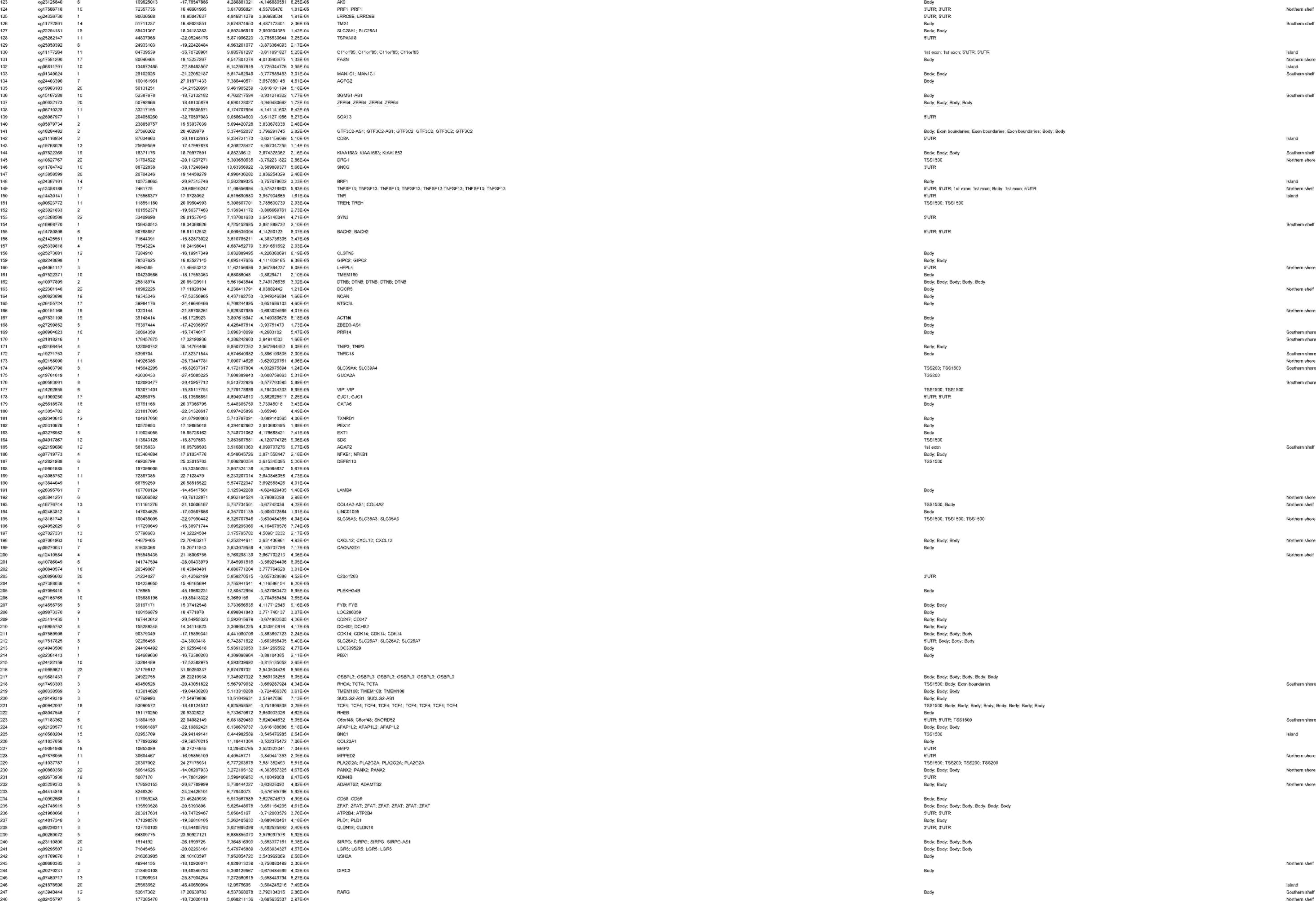
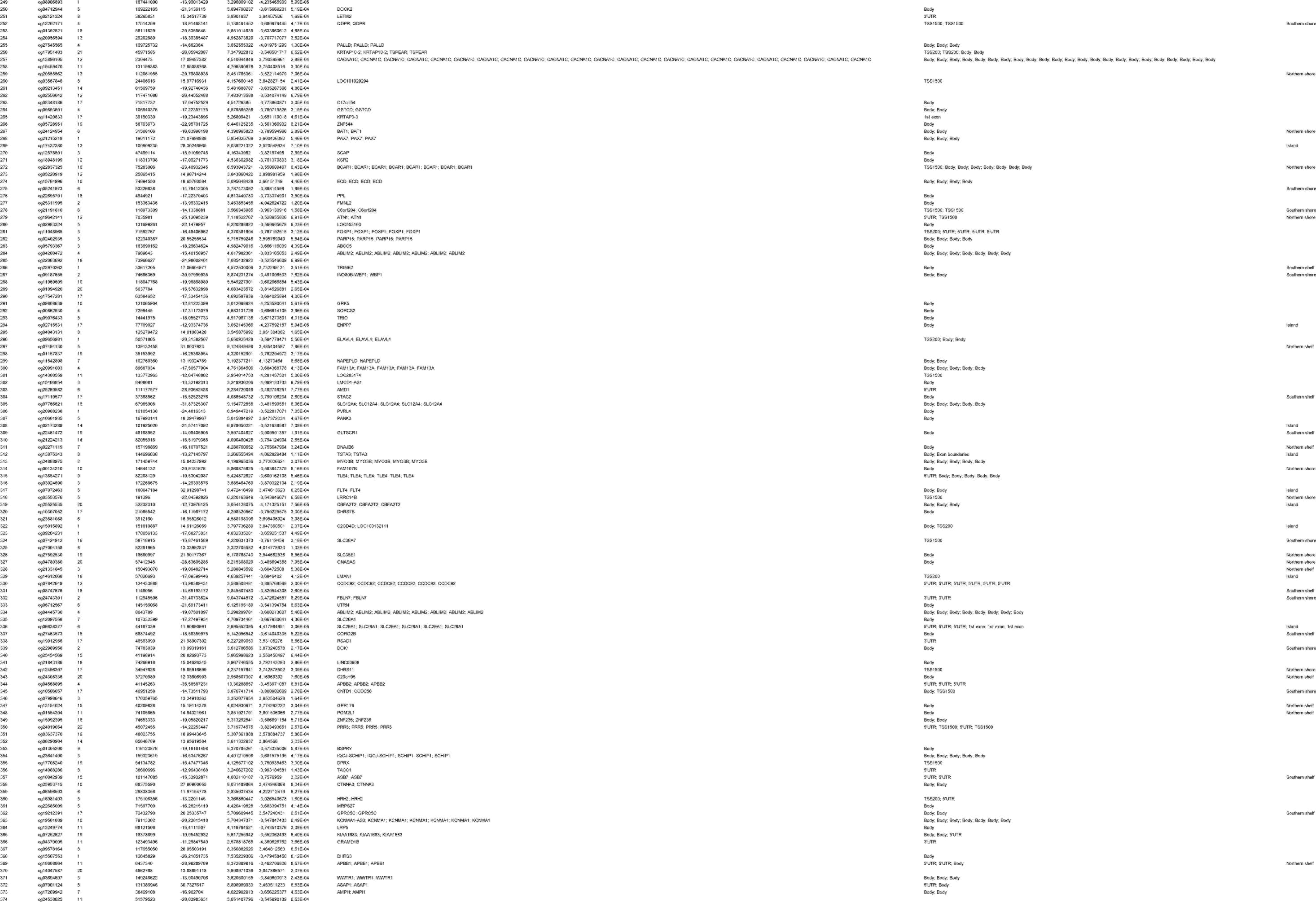
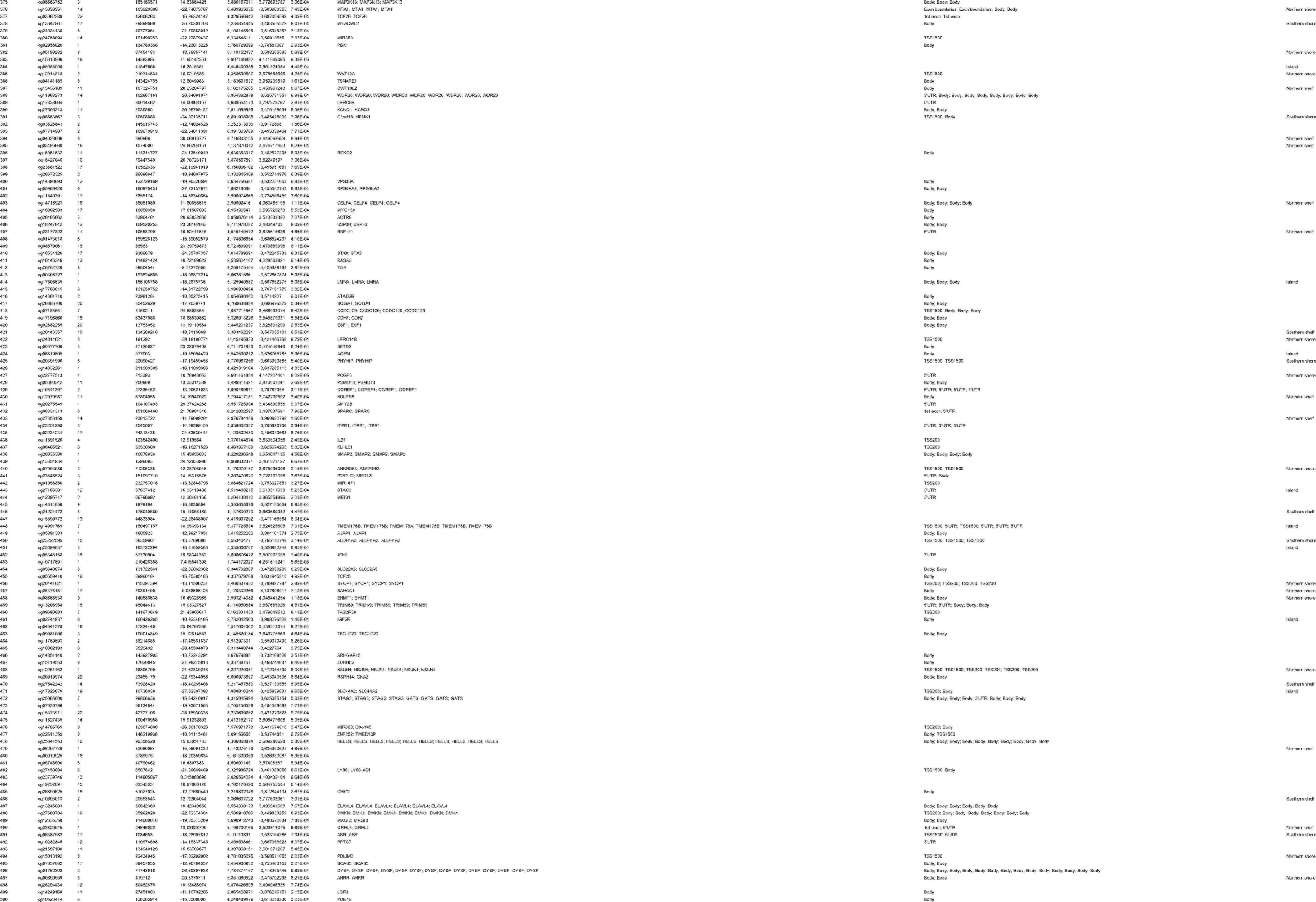
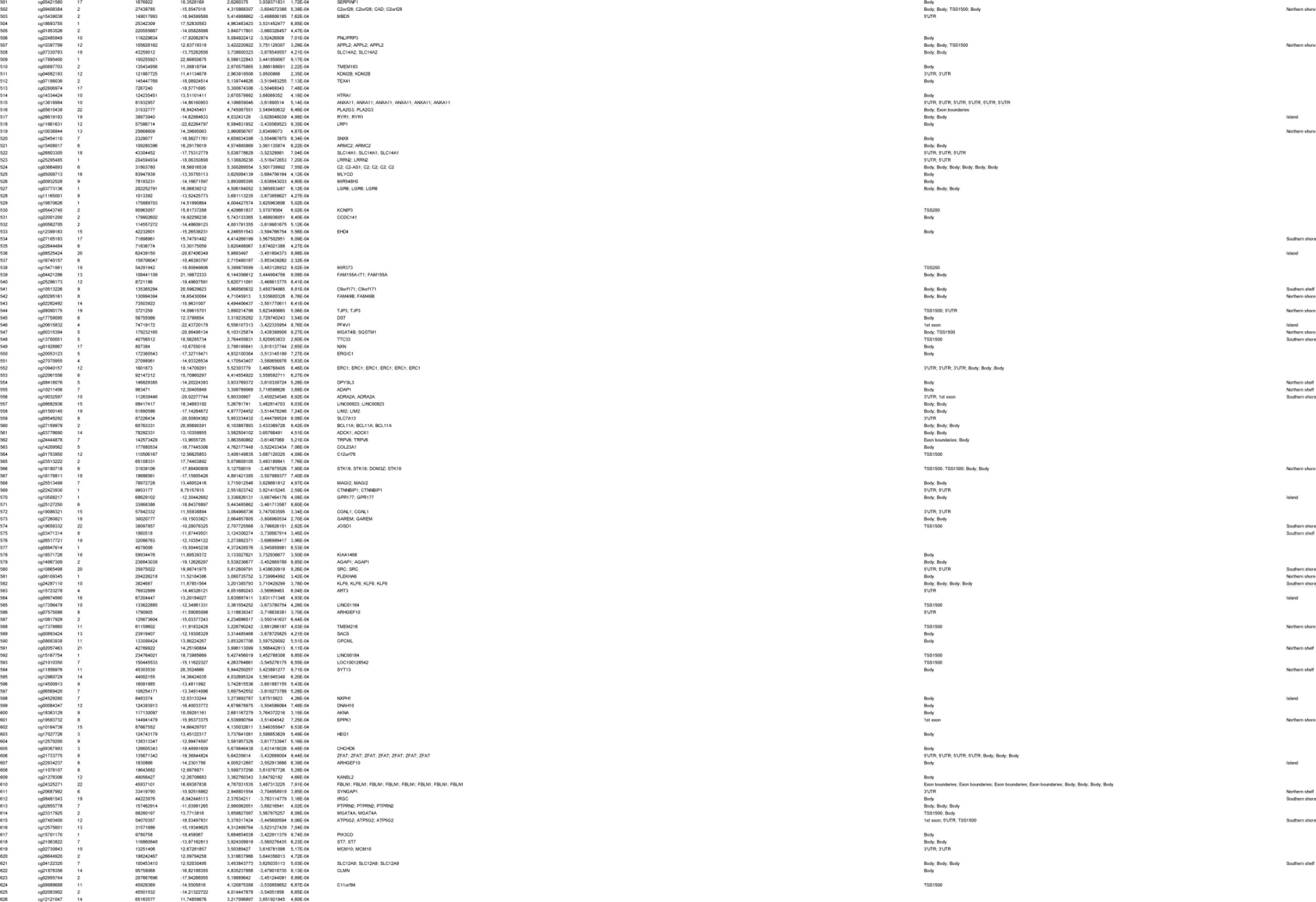
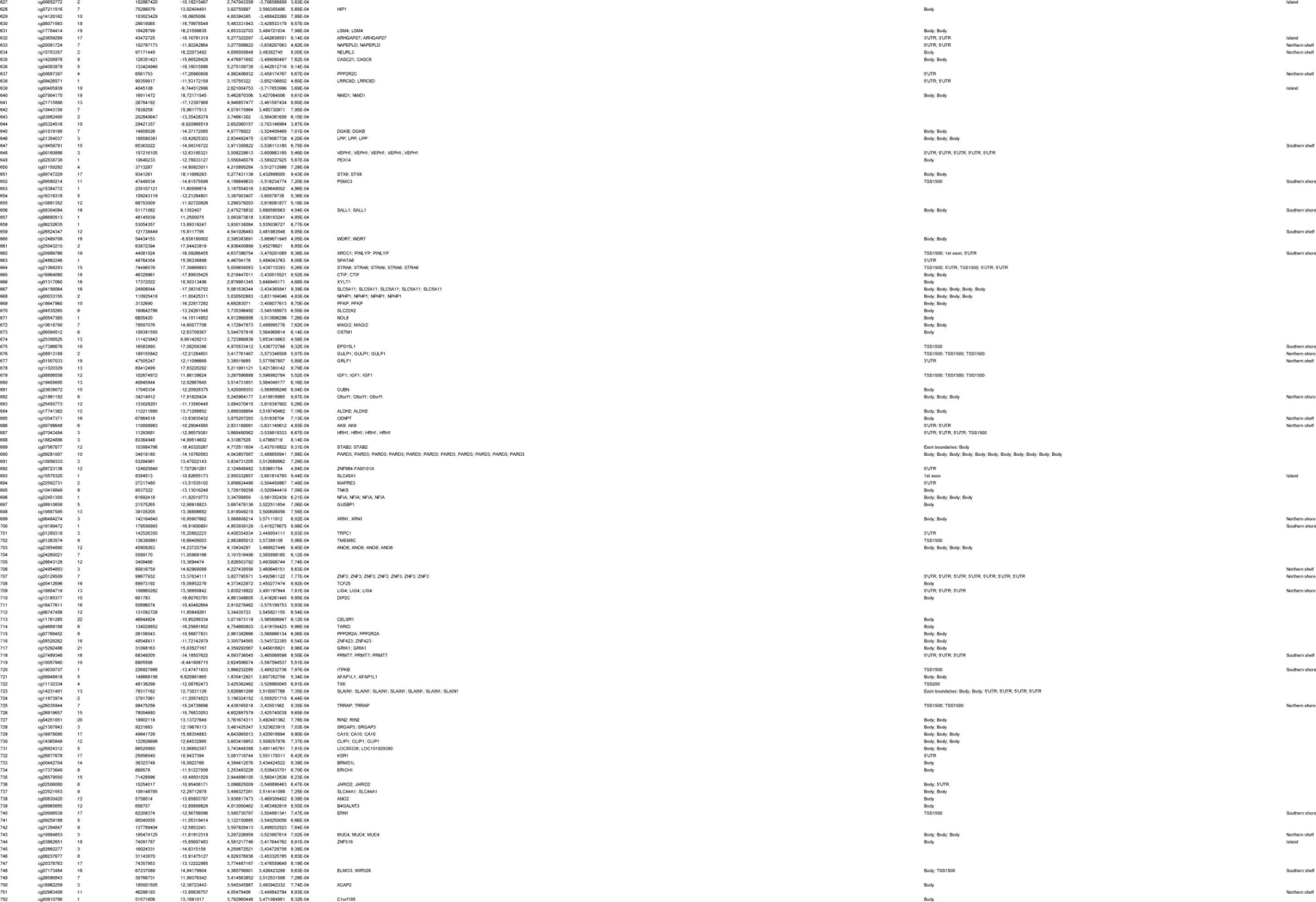
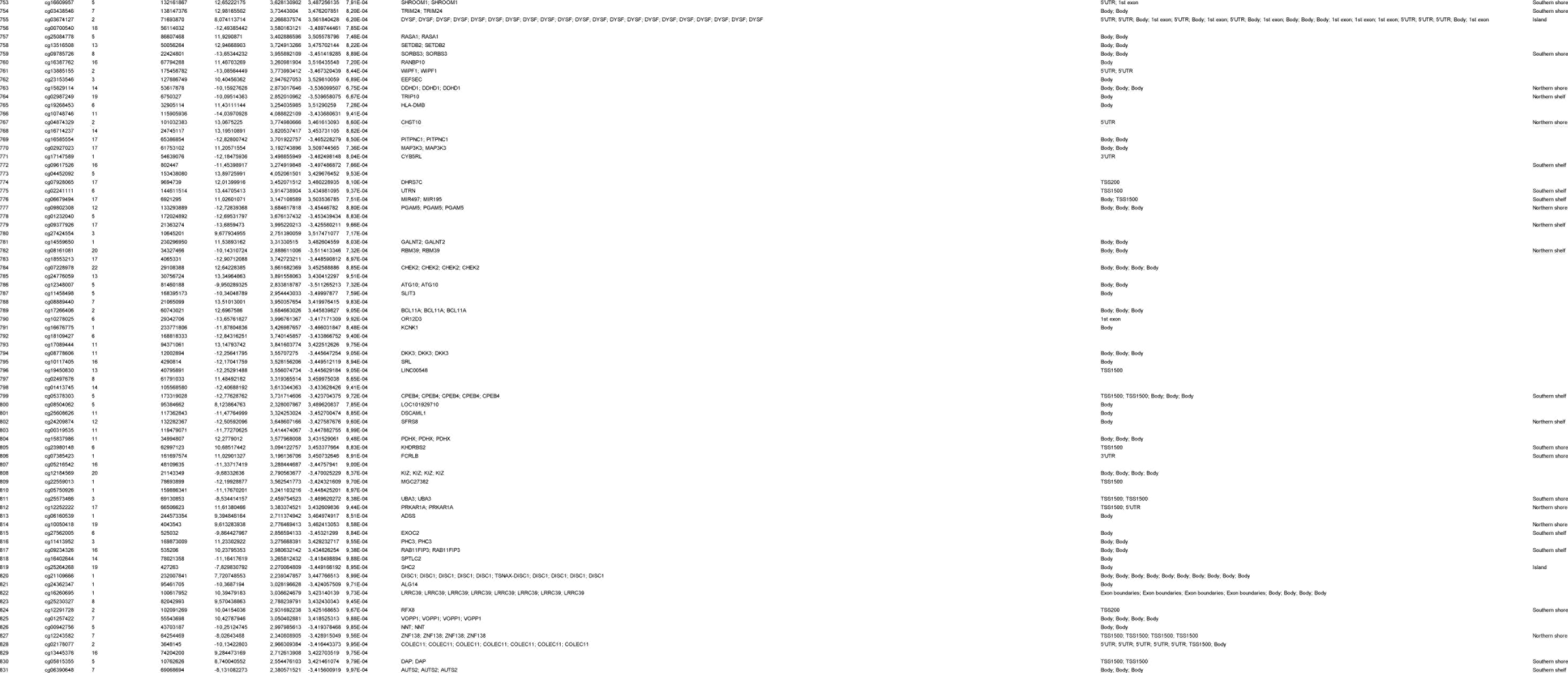
Top ranked differentially hydroxymethylated positions (DHPs) in the dorsal raphe nuclei (DRN) Displayed for each ranked probe is the chromosomal location and position (Ensembl GRCh37 assembly), the regression estimate (RE) for the Braak stage-associated analysis, the standard error (SE), the t-statistics, the accompanying p values, the Illumina gene annotation (UCSC Annotation), the gene feature (TSS1500, 200 to 1500 nucleotides (nt) upstream of transcription start site (TSS); TSS200, up to 200 nt upstream of TSS; 5′UTR, 5′untranslated region; Body, gene body; 3′UTR, 3′ untranslated region) and the cytosine-phosphate-guanine (CpG) island feature. Probes are ranked based on a combined p value (cut-off = p < 0.001) and regression estimate ranking.

**Supplementary Table 4.**
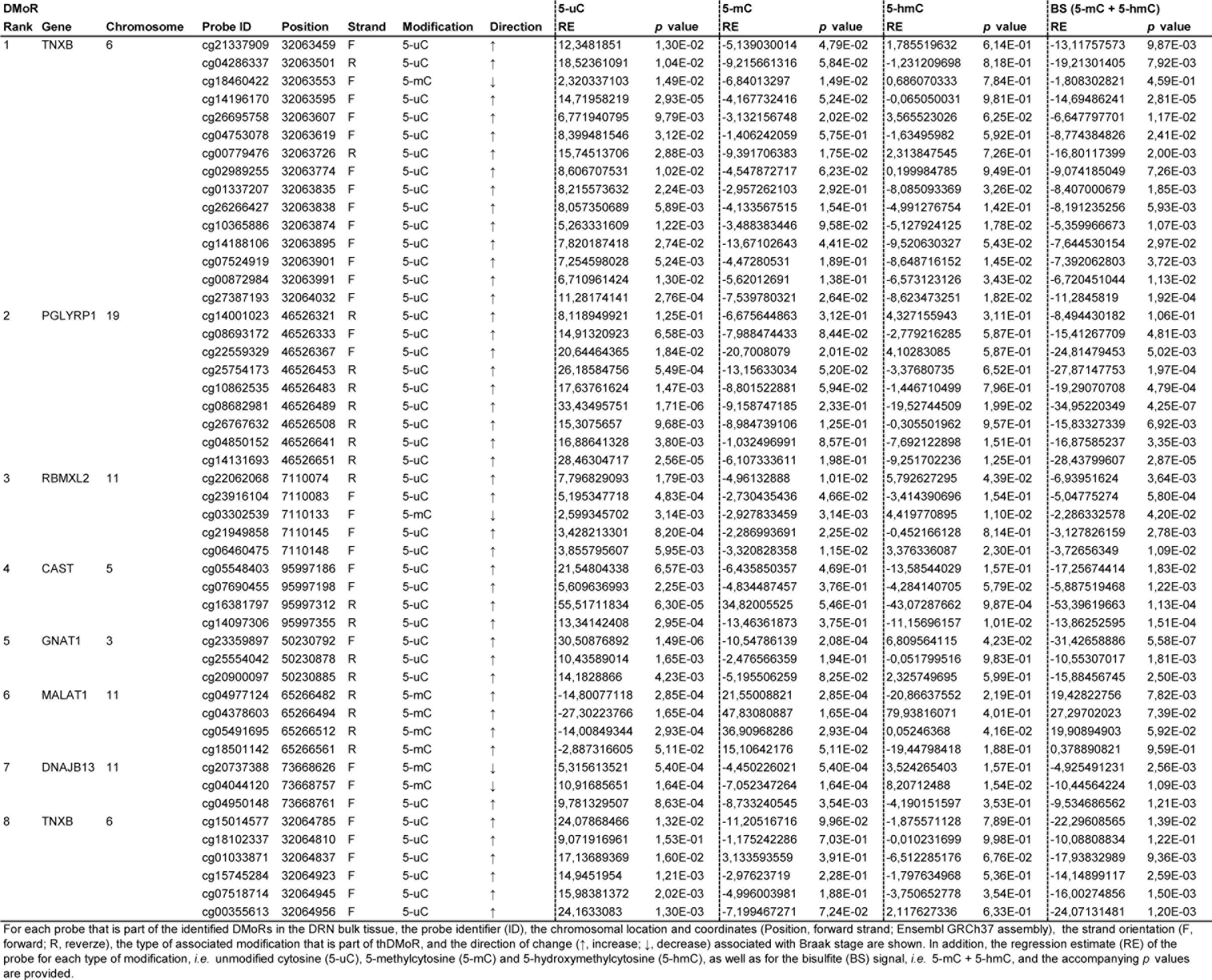
Overview of individual probes belonging to the differentially modified regions (DMoRs) in the dorsal raphe nucleus (DRN) bulk tissue.

**Supplementary Table 5.** Differentially unmodified-(DURs) and differentially methylated regions (DMRs) in the dorsal raphe nuclei (DRN) DURs (5-uC) Displayed are the differentially unmodifed regions (DURs) and the differentially methylated region (DMR) associated with Braak stage in the dorsal raphe nuclei (DRN). For each region, the Illumina gene annotation (UCSC Annotation), the chromosomal position and coordinates (Ensembl GRCh37 assembly), the number of probes, the *p* value, the multiple testing corrected *p* Šidák value, the gene feature (TSS, transcription start site; 5’UTR, 5’untranslated region; CDS, coding sequence; NC, non-coding), the cytosine-phosphate-guanine (CpG) island feature and the association with increasing Braak stage are shown. All regions are ranked based on their *p* Šidák value with *p* Šidák < 0.05 as a cut-off.

| Rank | USCS annotation | Chromosome | Start | End | Probes | *p* value | *p* Šidák value | Gene feature | CpG island feature | Association |
| --- | --- | --- | --- | --- | --- | --- | --- | --- | --- | --- |
| 1 | PGLYRP1 | 19 | 46526321 | 46526652 | 9 | 3,19E-13 | 7,24E-10 | Intergenic | Island | ↑ |
| 2 | TNXB | 6 | 32063607 | 32064033 | 11 | 2,63E-12 | 4,64E-09 | CDS | Island | ↑ |
| 3 | CAST | 5 | 95997186 | 95997356 | 4 | 3,35E-09 | 1,48E-05 | Intergenic | Shore | ↑ |
| 4 | GNAT1 | 3 | 50230792 | 50230886 | 3 | 1,14E-08 | 9,09E-05 | Intron; CDS | Island | ↑ |
| 5 | RBMXL2 | 11 | 7110074 | 7110149 | 5 | 1,07E-08 | 1,07E-04 | Intergenic | Island | ↑ |

**DMR (5-mC)**
| Rank | USCS annotation | Chromosome | Start | End | Probes | *p* value | *p* Šidák value | Gene feature | CpG Island feature | Association |
| --- | --- | --- | --- | --- | --- | --- | --- | --- | --- | --- |
| 1 | MALAT1 | 11 | 65266482 | 65266562 | 4 | 7,03E-09 | 6,61E-05 | NC exon | Shore | ↑ |

**Supplementary Table 6.**
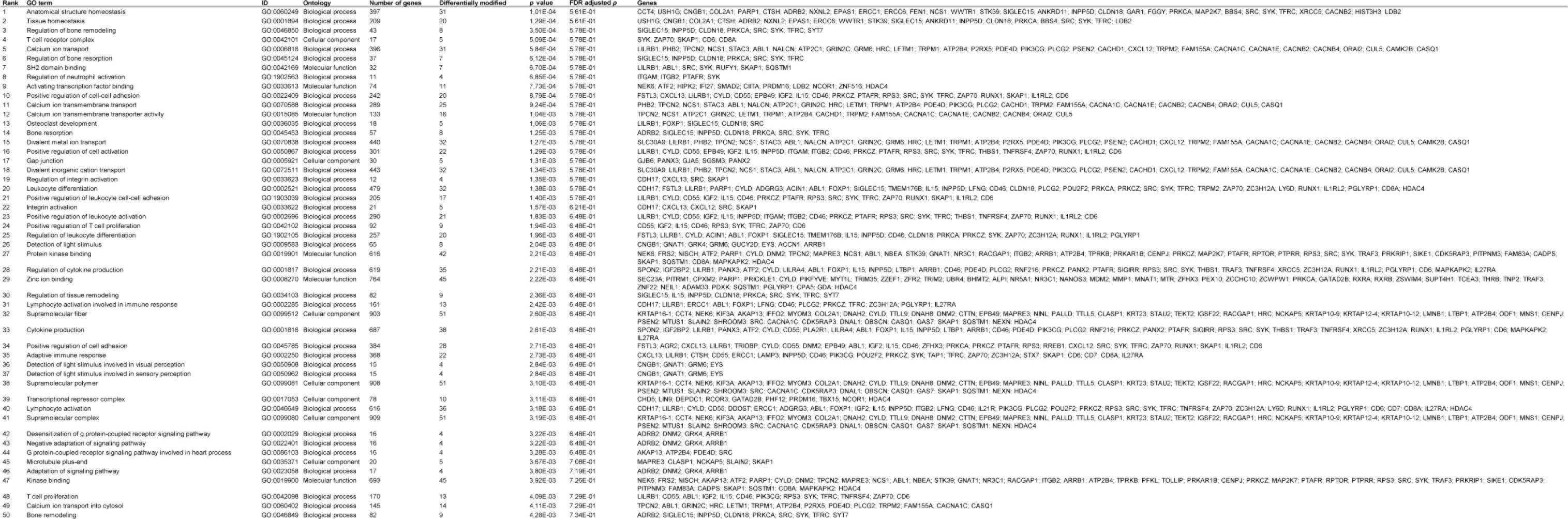
Top 50 enriched Gene Ontology (GO) terms for unmodified cytosine (5-uC) in the dorsal raphe nucleus (DRN) Displayed for each ranked GO term is the identifier (ID), the related ontology, the total number of genes belonging to the GO term, the number of genes that are differentially modified, the p value, the false discovery rate (FDR) adjusted p value and the names of the differentially modified genes. The GO terms are ranked based on their p value.

**Supplementary Table 7.**
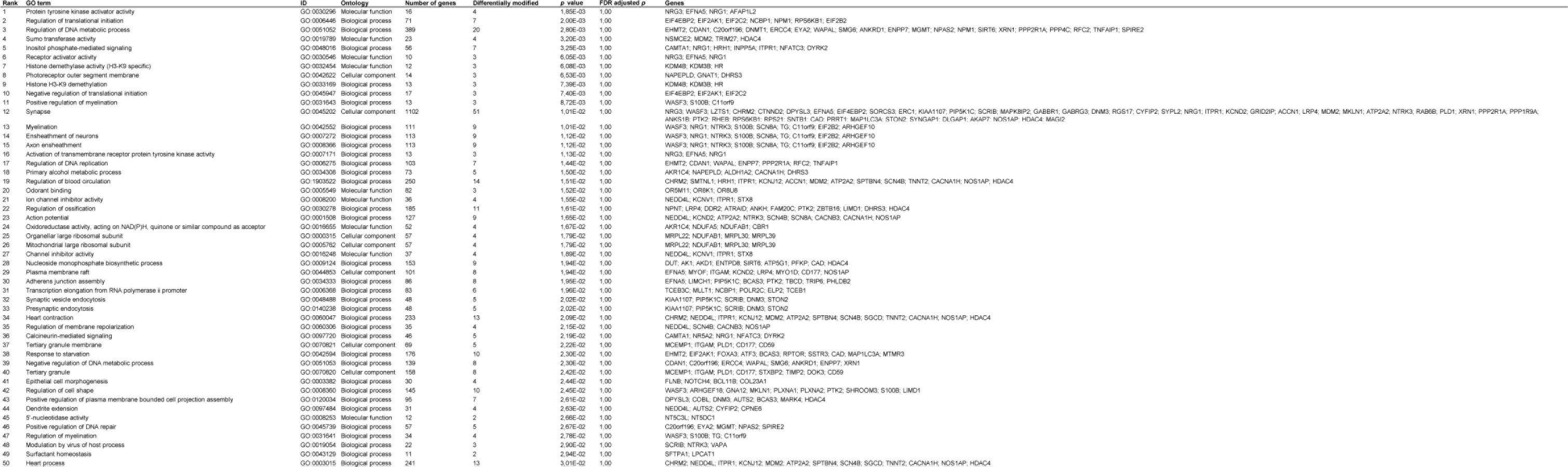
Top 50 enriched Gene Ontology (GO) terms for 5-methylcytosine (5-mC) in the dorsal raphe nucleus (DRN) Displayed for each ranked GO term is the identifier (ID), the related ontology, the total number of genes belonging to the GO term, the number of genes that are differentially modified, the p value, the false discovery rate (FDR) adjusted p value and the names of the differentially modified genes. The GO terms are ranked based on their p value.

**Supplementary Table 8.**
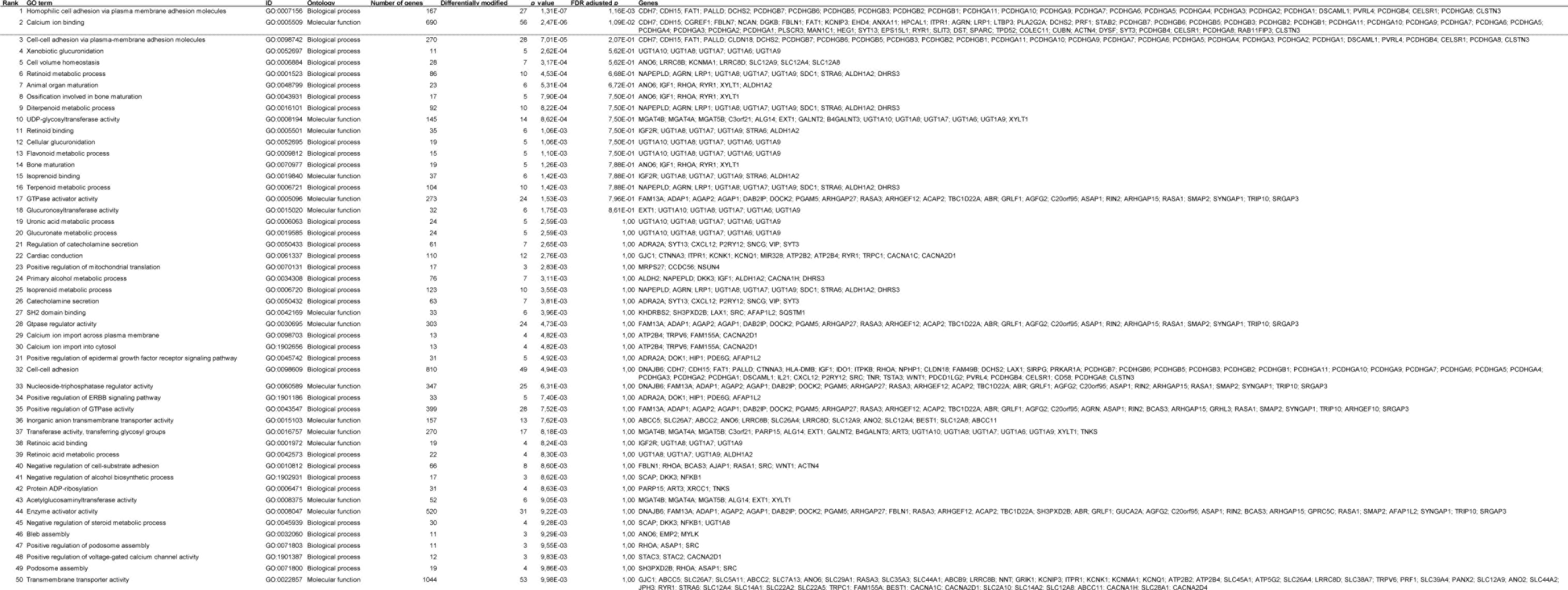
Top 50 enriched Gene Ontology (GO) terms for 5-hydroxymethylcytosine (5-hmC) in the dorsal raphe nucleus (DRN) Displayed for each ranked GO term is the identifier (ID), the related ontology, the total number of genes belonging to the GO term, the number of genes that are differentially modified, the p value, the false discovery rate (FDR) adjusted p value and the names of the differentially modified genes. The GO terms are ranked based on their p value and those that are significantly enriched after multiple testing correction (FDR adjusted p < 0.05) are seperated from the other terms by a dashed line.

**Supplementary Table 9.** Differentially bisulfite methylated regions (DbsMRs) in the dorsal raphe nucleus (DRN) bulk tissue.

| Rank | USCS annotation | Chromosome | Start | End | Num. of probes | *p* value | *p* Šidák value | Gene feature | CpG island feature |
| --- | --- | --- | --- | --- | --- | --- | --- | --- | --- |
| 1 | TNXB | 6 | 32063595 | 32064033 | 12 | 3,36E-12 | 5,71E-09 | CDS | Island |
| 2 | PGLYRP1 | 19 | 46526321 | 46526652 | 9 | 2,85E-15 | 6,55E-09 | Intergenic | Island |
| 3 | TNXB | 6 | 32064785 | 32064957 | 6 | 6,81E-05 | 2,98E-04 | CDS | Island |
| 4 | CAST | 5 | 95997186 | 95997356 | 4 | 3,94E-06 | 1,74E-02 | Intergenic | Shore |
| 5 | GNAT1 | 3 | 50230792 | 50230886 | 3 | 3,07E-06 | 2,46E-02 | Intron + CDS | Island |

For each DbsMR in the DRN bulk tissue, the Illumina gene annotation (UCSC Annotation), the chromosomal position and coordinates (Ensembl GRCh37 assembly), the number of probes, the *p* value, the multiple testing corrected *p* Šidák value, the gene feature (CDS, coding sequence) and the cytosine-phosphate-guanine (CpG) island feature are shown. All regions are ranked based on their *p* Šidák value with *p* Šidák < 0.05 as a cut-off.

**Supplementary Table 10.**
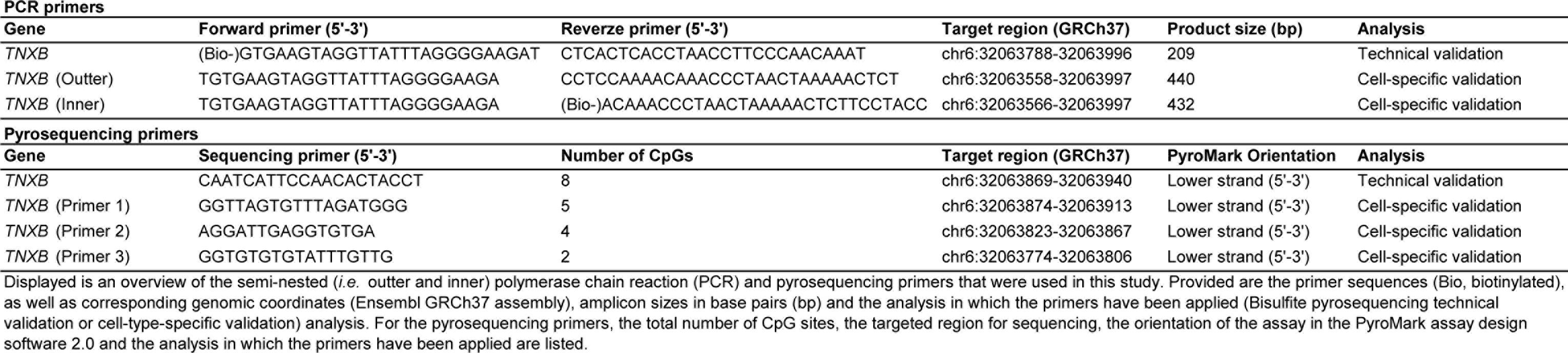
Polymerase chain reaction and pyrosequencing primers.

**Supplementary Table 11.** Overview of the limiting dilution bisulfite pyrosequencing (LDBSP) parameters.

|  | ***TNXB*** |  |  |
| --- | --- | --- | --- |
| *Singleplex PCR* | **Control** | **AD** | **Total** |
| **Number (+ percentage) of reactions with PCR product** | 104 (13.13) | 133 (16.79) | 237 (14.96) |
| *Allele estimation* |  |  |  |
| **Number (+ percentage) of reactions with 1 allele** | 97 (93.27) | 120 (90.23) | 217 (91.56) |
| **Number (+ percentage) of reactions with 2 alleles** | 5 (4.81) | 8 (6.02) | 13 (5.49) |
| **Number (+ percentage) of reactions with 3 alleles** | 2 (1.92) | 5 (3.76) | 7 (2.95) |
| **Number (+ percentage) of multi-allele reactions** | 7 (6.73) | 13 (9.77) | 20 (8.44) |
| **Number (+ percentage) of recovered alleles** | 113 (3.14) | 151 (4.19) | 264 (3.67) |

A summary of all the LDBSP parameters for pools of 50 serotonergic neurons isolated from unfixed post-mortem brain tissue using laser capture microdissection (LCM). Displayed are the total number of obtained polymerase chain reaction (PCR) products per experimental groups and the total amount of estimated alleles. Please refer to the written text for further specifications on the allele estimation method and criteria.

